# SATB2 induction of a neural crest mesenchyme-like program drives invasion and drug resistance in melanoma

**DOI:** 10.1101/2020.11.01.364406

**Authors:** Maurizio Fazio, Ellen van Rooijen, Michelle Dang, Glenn van de Hoek, Julien Ablain, Jeffrey K. Mito, Song Yang, Andrew Thomas, John Michael, Tania Fabo, Rodsy Modhurima, Patrizia Pessina, Charles Kaufman, Yi Zhou, Richard M. White, Leonard I. Zon

## Abstract

Recent genomic and scRNA-seq analyses of melanoma identified common transcriptional states correlating with invasion or drug resistance, but failed to find recurrent drivers of metastasis. To test whether transcriptional adaptation can drive melanoma progression, we made use of a zebrafish *mitfa:BRAFV600E;tp53-/-* model, in which malignant progression is characterized by minimal genetic evolution. We undertook an overexpression-screen of 80 epigenetic/transcriptional regulators and found neural crest-mesenchyme developmental regulator SATB2 to accelerate aggressive melanoma development. Its overexpression induces invadopodia formation and invasion in zebrafish tumors and human melanoma cell lines. SATB2 binds and activates neural crest-regulators, including *pdgfab* and *snai2.* The transcriptional program induced by SATB2 overlaps with known MITF^low^AXL^hlgh^ and AQP1^+^NGFR1^high^ drug resistant states and functionally drives enhanced tumor propagation and resistance to Vemurafenib *in vivo.* Here we show that melanoma transcriptional rewiring by SATB2 to a neural crest mesenchyme-like program can drive invasion and drug resistance in endogenous tumors.

---

Both sequencing of patient precursor lesions (Shain et al., 2015) and modeling in transgenic animals (Perez-Guijarro, Day, Merlino, & Zaidi, 2017; van Rooijen, Fazio, & Zon, 2017) showed that acquisition of oncogenic *BRAF* or *NRAS* mutations alone is insufficient to drive melanoma initiation. Reconstruction of the clonal history of advanced human melanoma suggests that few driver mutations, including loss of tumor suppressors, are acquired in early transformed melanocytes, and that high mutational burden and genomic instability are later events in malignant progression (Birkeland et al., 2018; Ding et al., 2014). A pan-cancer whole-genome analysis of metastatic solid tumors, including 248 melanoma patients, showed a surprisingly high degree of similarity in mutational landscape and driver genes between metastatic tumors and their respective primaries, as well as across multiple metastatic lesions within the same patient (Priestley et al., 2019). The lack of recurrently mutated genes in metastasis, and the limited genetic diversity across metastatic sites suggests that genetic selection of mutated drivers is unlikely to be responsible for metastatic progression. Despite the high degree of mutational burden and genetic intra- and inter-patient heterogeneity observed in melanoma, several bulk RNA-seq and scRNA-seq analyses of metastatic melanoma patient samples have identified common recurrent transcriptional states correlating with invasion (mesenchymal signature) (Verfaillie et al., 2015), drug resistance (MITF^low^/AXL^high^) (Tirosh et al., 2016), a neural crest-like state correlating with minimal residual disease persistence during targeted therapy with MAPK inhibitors (MITF^low^/NGFR1^high^/AQP1^high^) (Rambow et al., 2018), and even response to immunotherapy (NGFR1^high^) (Boshuizen et al., 2020). The existing literature on these often less proliferative cell states has been limited to *in vitro* perturbations, or relied on transplant models, and as such their role in oncogenesis outside of iatrogenic drug treatment remains less clear.

Using a zebrafish transgenic melanoma model, we have previously shown that in a cancerized field of melanocytes carrying driving oncogenic mutations, such as *BRAF^V600E^* and loss of *tp53,* only a few cells or subclones will go on to develop a malignant tumor, and that this event is marked by re-expression of developmental NC-markers *sox10* and *crestin* (Kaufman et al., 2016; McConnell et al., 2018). Similar to the observation in human metastatic patient samples, malignant progression from transformed melanocytes in experimental animal models is not explained by the very limited genetic evolution observed in these tumors (Yen et al., 2013). Taken together, these observations in human patients and animal disease models raise the question whether epigenetic adaptation, rather than genetic selection, can contribute to tumor progression by altering the transcriptional state of the tumor.

Here, we address this hypothesis by high throughput genetic perturbation of 80 epigenetic/transcriptional regulators in endogenous primary tumors, by leveraging a transgenic zebrafish melanoma model uniquely suited to precisely and rapidly perturbate tumor development *in vivo* (Ceol et al., 2011), at a scale and speed much greater than possible with genetically engineered murine melanoma models. We identify Special AT-rich Binding protein 2 (SATB2) as a novel accelerator of melanoma onset driving an aggressive phenotype in primary tumors suggestive of metastatic spreading. SATB2 acts as a transcriptional regulator by recruiting members of the Acetyl Transferase and Histone demethylase complexes to target genes and altering the local chromatin organization and activation state (L. Q. Zhou et al., 2012). Its expression has been shown to correlate with patient outcome in several tumor types (Naik & Galande, 2019; Yu, Ma, Shankar, & Srivastava, 2017). In different tissues and tumor types SATB2 has been shown to play a role in oncogenic transformation and proliferation, or Epithelial-to-Mesenchymal Transition (EMT), migration, and self-renewal (Gan et al., 2017; Naik & Galande, 2019; Nayak et al., 2019; Wang et al., 2019; Wu et al., 2016; Xu et al., 2017; Yu, Ma, Ochoa, Shankar, & Srivastava, 2017)). SATB2 is a transcription factor and chromatin remodeler with a well-conserved structure and expression pattern across chicken, mouse and zebrafish during the development and migration of the cranial neural crest (CNC) and neuronal development. It is required for the development of the exo-mesenchymal lineages of the CNC (Sheehan-Rooney, Palinkasova, Eberhart, & Dixon, 2010; Sheehan-Rooney, Swartz, Lovely, Dixon, & Eberhart, 2013), and neuronal axon formation (McKenna et al., 2015; Shinmyo & Kawasaki, 2017). In facts, SATB2 inactivating mutations in humans have been associated with cleft palate, intellectual disability, facial dysmorphism, and development of odontomas, defining a neurocristopathy referred to as SATB2-associated syndrome (Kikuiri et al., 2018; Zarate et al., 2019; Zarate & Fish, 2017). Through a combination of zebrafish *in vivo* allotransplants and validation in human melanoma cell lines, we show that SATB2 drives enhanced invasion via invadopodia formation and an EMT-like phenotype. Mechanistically, chromatin and transcriptional characterization of primary zebrafish SATB2 tumors vs. EGFP controls via ChIP-seq and RNA-seq shows SATB2 to bind and induce transcriptional activation of neural crest regulators, including *snai2* and *pdgfab.* The transcriptional program induced by SATB2 overexpression is conserved between zebrafish and human melanoma, and overlaps with the aforementioned MITF^low^/AXL^high^ (Tirosh et al., 2016) and neural crest-like MITF^low^/NGFR1^high/^AQP1^high^ drug resistant states (Rambow et al., 2018). Finally, we show SATB2 transcriptional rewiring to functionally drive enhanced tumor propagation and resistance to MAPK inhibition by Vemurafenib in zebrafish tumor allografts *in vivo.*

## Results

### In vivo overexpression screen of epigenetic factors identifies SATB2 as melanoma accelerator

To interrogate whether epigenetic reprogramming can accelerate melanoma development, we utilized a genetic discovery driven approach and undertook an *in vivo* overexpression screen. We tested 80 chromatin factors: 15 pools of 5 factors, and 6 additional single factors including previously published positive controls SETDB1, SUV39H1 (Ceol et al., 2011) (see Supplementary Table 1). EGFP was used as a negative control, and we tested CCND1 as an additional positive control (known driver often amplified in melanoma (Cancer Genome Atlas, 2015)). As a screening platform, we leveraged a zebrafish melanoma model driven by tissue-specific expression of human oncogenic *BRAF^V600E^* in a *tp53* and *mitfa*-deficient background (Ceol et al., 2011). *Tg(mitfa:BRAF^V600E^); tp53-/-; mitfa-/-* zebrafish lack melanocytes and do not develop melanoma. Mosaic integration of the transposon-based expression vector MiniCoopR (MCR), rescues melanocyte development by restoring *mitfa,* while simultaneously driving tissuespecific expression of candidate genes (Fig. 1A). Thus in mosaic F0 transgenics all melanocytes express the candidate gene tested (described in Ceol et al. (Ceol et al., 2011)). We identified 6 candidate pools, of which Pool F (Figure 1B) had the strongest acceleration. Single factor validation of Pool F (Figure 1C) showed that SATB2-overexpression (Supplemental Figure 1A-B) is sufficient to accelerate melanoma development (median onset of 12 weeks, compared to 21.4 weeks for MCR:EGFP) (Figure 1C). MCR:SATB2 tumors appear grossly aggressive (Figure 1D, Supplemental Figure 1C, Supplemental Video 1), are invasive, and melanoma cells are frequently observed in internal organs and spreading along the spinal cord, while organ involvement is rarely observed in MCR:EGFP (Figure 1E-G, Supplemental Figure 1C-F). SATB2 melanoma development is *BRAFV600E* and *tp53-/-*dependent (Supplemental Figure 2A) and tissue-specific CRISPR/CAS9-induced loss of zebrafish *satb2* did not affect tumor onset compared to control CRISPR targeting *tp53* (Ablain et al., 2018)(Supplemental Figure 2B-C). The acceleration phenotype is not due to increased cellular proliferation, since SATB2-overexpression in primary zebrafish tumors (Supplemental Figure 3A) or in a panel of human melanoma cell lines via a TETon tetracycline inducible lentiviral vector (here referred to as iSATB2) did not result in increased proliferation (Supplemental Figure 3B,C). Our data demonstrate that SATB2-overexpression accelerates melanoma malignant progression without affecting proliferation, but SATB2 is neither necessary nor sufficient for melanoma initiation.

**Figure 1:**
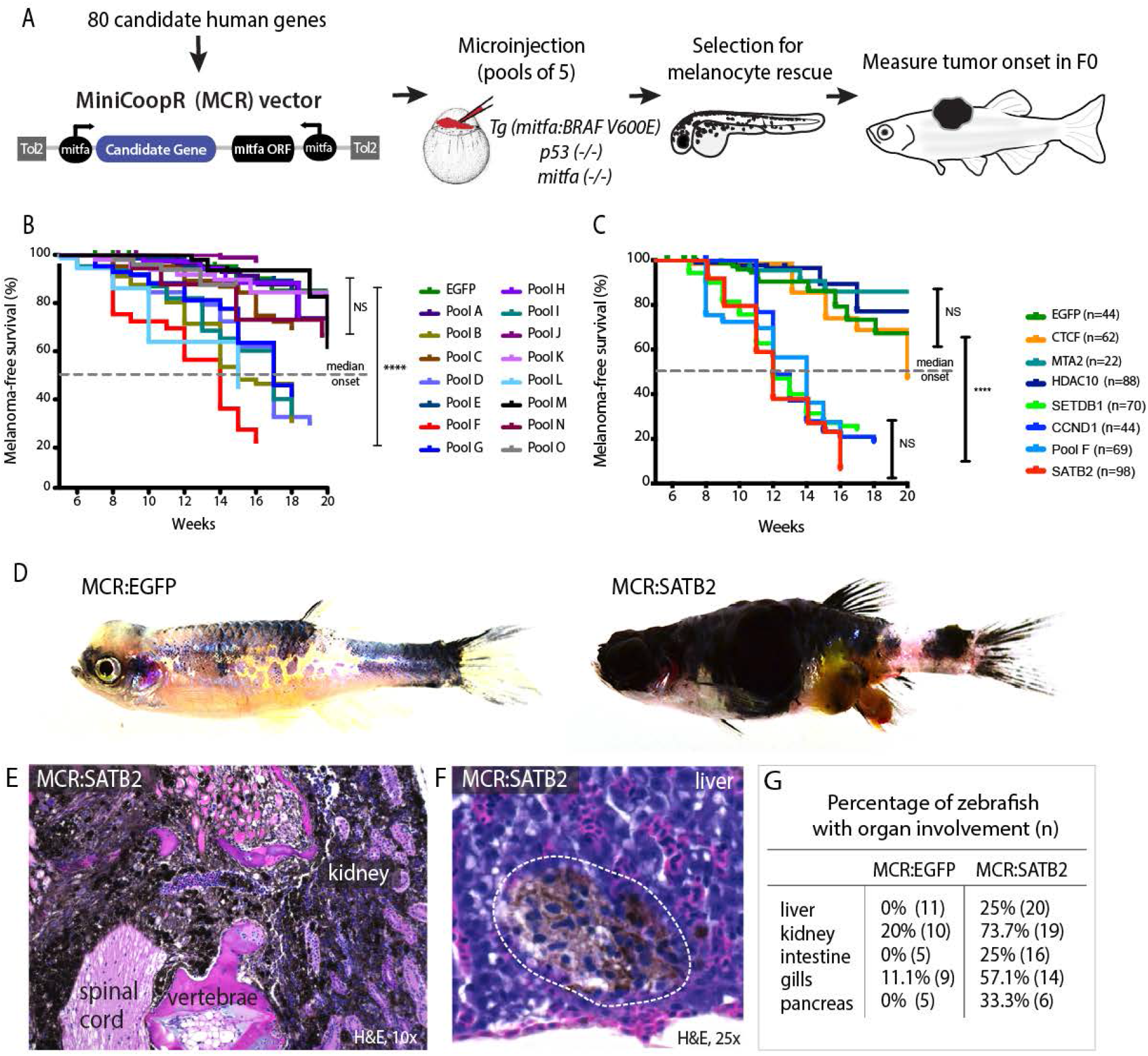
Overexpression screen of epigenetic regulators identifies SATB2 as an accelerator of melanoma formation in zebrafish. (A) Schematic overview of screening strategy: MCR expression vector-based reintroduction of the melanocyte master regulator *mitfa,* rescues melanocytes and melanoma development. All rescued melanocytes in F0 microinjected embryos also express a candidate human factor. (B) Kaplan-Meier melanoma-free survival curves of pooled chromatin factor screen. Six pools were significant (logrank, *p*<0.0001****). Pool F (red) had the strongest acceleration effect (Median onset 14 weeks). (C) Single factor validation of pool F identifies SATB2 to induce accelerated melanoma onset (Median onset 12 weeks, logrank, *p*<0.0001****). (D) Zebrafish MCR:SATB2 tumors are aggressive compared to an MCR:EGFP age-matched control. (E) Histopathological analysis revealed that MCR:SATB2 tumors are highly invasive, here shown to invade through the spinal cord, vertebrae and kidney. (F) Isolated melanoma cell clusters were found in the liver, and (G) frequent organ involvement is observed in MCR:SATB2 compared to MCR:EGFP controls.

### SATB2 overexpression leads to invadopodia formation and increases migration and invasion in vitro and in vivo

Given the lack of proliferation changes, we next asked whether SATB2’s aggressive tumor phenotype and internal tumors might be explained by an increase in tumor migration and/or invasion. Using primary zebrafish melanoma *in vitro* cell cultures we performed scratch migration assays, that confirmed a heightened migration potential of 3 independent MCR:SATB2 cell lines compared to 3 independent MCR:EGFP cell lines (Supplementary Figure 4A). Cytoskeletal staining of zebrafish melanoma *in vitro* cultures (Zmel1 MCR:EGFP and 45-3 MCR:SATB2) with phalloidin revealed the presence of strong F-actin positive foci in MCR:SATB2 cells (Figure 2A), reminiscent of invadopodia. Invadopodia are cell protrusions involved in metastatic spreading by facilitating anchorage of cells to, and local degradation of the ECM (Murphy & Courtneidge, 2011), and are known to be regulated by the PDGF-SRC pathway and EMT. During normal development, invadopodia-like physiologically equivalent structures called podosomes are utilized by neural crest cells to migrate (Murphy & Courtneidge, 2011; Murphy et al., 2011). Co-localization of F-actin with invadopodium structural component Cortactin in MCR:SATB2 melanoma cell lines confirmed these foci to be invadopodia (Figure 2A and Supplementary Figure 4B). Much like *in vitro* cultures, primary MCR:SATB2 tumors also showed abundant Cortactin expression compared to MCR:EGFP (Figure 2B). Proteolytic activity of invadopodia induces local degradation of the extracellular matrix. To test whether SATB2 expressing cells form functional invadopodia, we plated cells onto Oregon green 488-conjungated gelatin-coated coverslips and assayed for matrix degradation 24-25 hours postseeding (Martin et al., 2012). MCR:SATB2 melanoma cell lines strongly degraded the gelatin matrix, with 49.2-57.9% of cells having degraded gelatin, compared to 5.4% of MCR:Empty control cells (Figure 2C, Supplementary Figure 4C-D). To validate the effect of SATB2 as a regulator of invadopodia formation in human melanoma, we utilized the panel of iSATB2 human melanoma cell lines. Upon SATB2 induction (Supplemental Figure 3B), all human melanoma cell lines robustly formed invadopodia and showed significantly increased matrix degradation (Figure 2D-E, Supplemental Video 2). In SKMEL2, a ten-fold induction of cells with matrix degradation was observed, from a baseline of 5.4%±4.5% (SD) of cells with matrix degradation to 58.2%±13.2% (SD) of cells after doxycycline treatment (Figure 2D-E), with evidence of F-actin and Cortactin co-localization in punctae below the cell nucleus, characteristic of invadopodia (Figure 2E, Supplementary Figure 4E).

**Figure 2:**
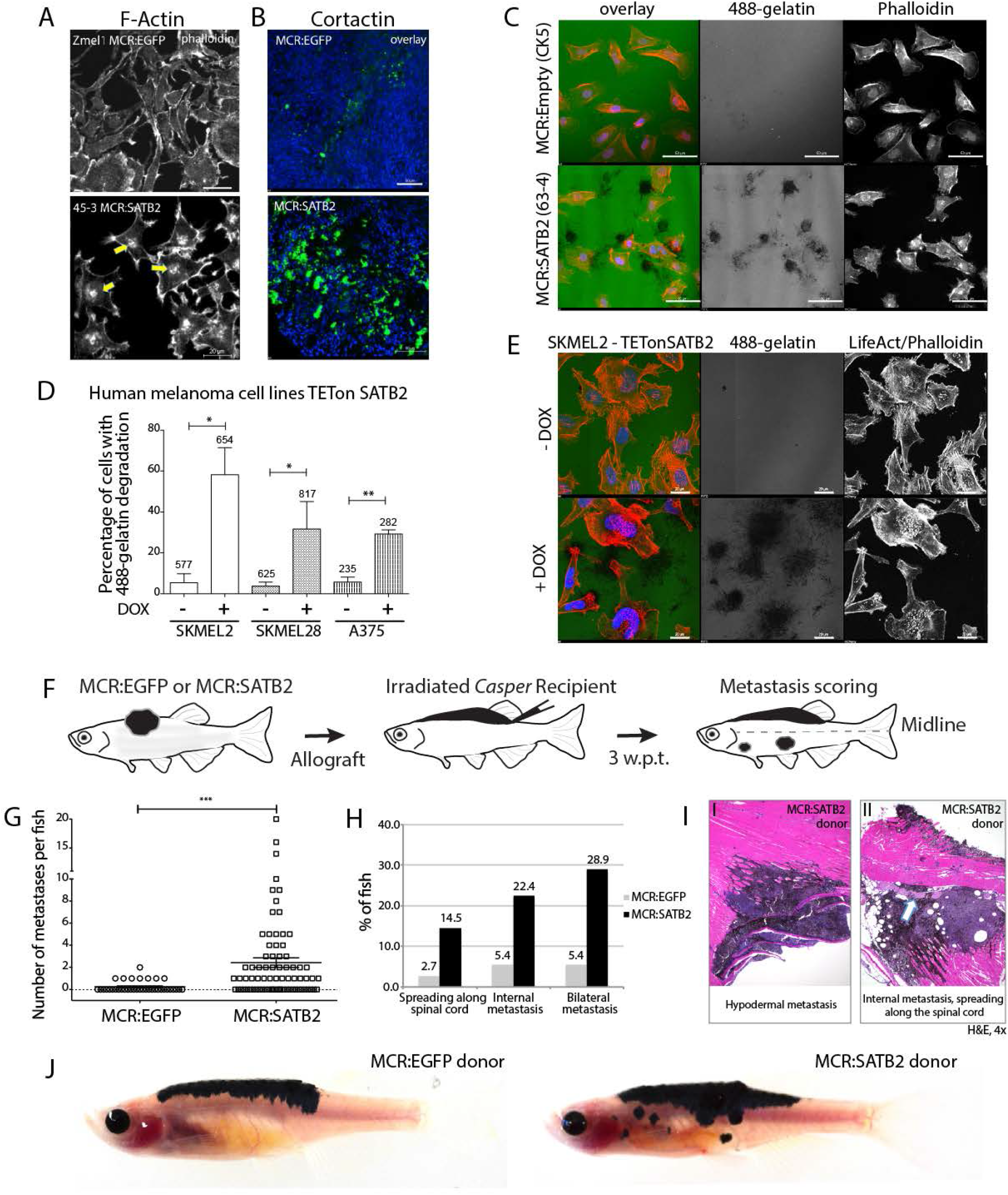
SATB2 leads to invadopodia formation and increased migration potential *in vitro* and *in vivo*. (A) Phalloidin staining for F-Actin in primary zebrafish melanoma cell culture, reveals the presence of F-actin positive foci in MCR:SATB2 (45-3) cells, which are not present in MCR:EGFP (zmel1) cells. Scale bar is 20 μm. (B) Primary tumor immunohistochemistry shows MCR:SATB2 tumors abundantly express Cortactin. Scale bar 50 μm. (C) MCR:SATB2 (63-4) cells show increased Oregon green 488-conjungated gelatin degradation, compared to MCR (CK5) cells 24-25 hours post-seeding. Scale bar is 50 μm. (D) Percentage of cells with degraded gelatin after SATB2 induction in iSATB2 human melanoma cell lines A375, SKMEL2 and SKMEL28 transduced with pInducer20- SATB2.Cells were induced in media +/− doxycycline for 48 hrs, and next seeded onto Oregon Green-gelatin in media +/− doxycycline for 24-25 hrs to assess matrix degradation. (E) Upon SATB2 induction in human melanoma cell line SKMEL2, cells form invadopodia and show increased matrix degradation. Scale bar is 20 μm. (F) Orthotropic allograft migration assay in transparent *casper* zebrafish. 300,000 primary pigmented primary melanoma cells were transplanted into the dorsum of irradiated *casper* recipients, which are monitored for the formation of pigmented distant metastases that have spread past the anatomical midline. Metastases are represented as black circles. (G) At the experimental end point at 3.5 weeks post-transplantation, 59.4%±2.3% (SEM; n=76, 7 donor tumors) of MCR:SATB2 transplants formed distant metastasis, compared to 21.8 %±4.5% (SEM; n=37, 5 donor tumors) of MCR:EGFP control transplants (*p*<0.0001). MCR:EGFP transplants developed an average 1.1±0.4 (SD) distant metastasis per fish, versus 4±4.2 (SD) in MCR:SATB2, where a maximum of 20 metastases per fish was observed. (H) MCR:SATB2 recipients more frequently developed bilateral (28.9%), and internal metastases (22.4%) compared to MCR:EGFP donors (5.4%), showing spreading along the neural tube (14.5% versus 2.7%). (I) Histopathology of MCR:SATB2 *casper* recipients showing a (I-I) hypodermal metastasis, and (I-II) internal metastasis with spreading along neural tube at 3.5 weeks-post transplantation.

To further validate whether the internal organ involvement in MCR:SATB2 primary tumors was due to an increased migratory potential *in vivo,* we used an orthotopic allotransplantation model in sub-lethally irradiated transparent *casper* zebrafish (Heilmann et al., 2015; P. Li, White, & Zon, 2011; White et al., 2008). First, we allotransplanted zebrafish melanoma-derived cell lines Zmel1 (MCR:EGFP) vs. 45-3 (MCR:SATB2) into irradiated *casper* recipients. This showed SATB2 to cause an invasive histological phenotype and reduction in the recipient’s overall survival compared to EGFP (45-3; n=31, median survival=25 days vs. Zmel1; n=31, median onset not reached at the experimental end point) (Supplemental Figure 5A-C). We then tested whether this difference was also present in primary tumors using an established *in vivo* migration assay (Heilmann et al., 2015) where we transplanted 300,000 primary pigmented melanoma cells into *casper* recipients, and monitored the formation of distant metastasis (Figure 2F). One week post-transplantation, 18.6% (11/59) of MCR:SATB2 recipients already developed metastases while none were observed in MCR:EGFP recipients (0/27) (Supplemental Figure 5D). At the experimental end point at 3-3.5 weeks-post transplantation, 59.4%±2.3% (SEM; n=76, 7 donor tumors) of MCR:SATB2 transplants formed distant metastasis, compared to 21.8%±4.5% (SEM; n=37, 5 donor tumors) of EGFP-control transplants (*p*<0.0001) (Figure 2GJ, and Supplemental Videos 3-4). In *casper* recipients that formed metastases, MCR:EGFP transplants developed an average of 1.1±0.4 (SD) distant metastases per fish, versus 4±4.2 (SD) in MCR:SATB2, where a maximum of 20 metastases per fish was observed (Figure 2G) (*p*<0.0001). Metastases most commonly spread to hypodermal sites (Figure 2H-J), similar to *in-transit* metastases that occur in human melanoma. MCR:SATB2 recipients more frequently developed bilateral metastases (28.9% versus 5.4% in MCR:EGFP), and internal metastases (22.4% versus 5.4%) which regularly spread along the neural tube (14.5% versus 2.7%) (Figure 2H-I). Collectively, these *in vitro* and *in vivo* data suggest that SATB2-overexpression induces invadopodia formation, increased migration and metastasis. SATB2 is endogenously expressed in human melanoma at the RNA (Supplementary Figure 6A) and protein level (Supplementary Figure 6B). Human melanoma patient genomic datasets publicly available on cBio portal show SATB2 to be infrequently but recurrently amplified in ~4-8% of patients (Supplementary Figure 6C) in 3 independent datasets of metastatic melanoma(Hugo et al., 2016; Snyder et al., 2014; Van Allen et al., 2015), and the high mRNA expression level correlate with poor survival in two independent metastatic melanoma patient datasets available on TIDE portal (GSE22153 and GSE8401, Supplementary Figure 6D).

### SATB2 binds and transcriptionally regulates neural crest- and EMT-associated loci

To gain insight into the mechanism underlying the SATB2-overexpression phenotype in melanoma, we performed ChIP-seq and RNA-seq on primary zebrafish tumors. We conducted ChIP-seq on MCR:SATB2 primary zebrafish melanomas to identify SATB2-bound target genes (antibody validation in Supplementary Figure 7A). HOMER motif analysis of ChIP-seq significant peaks for de novo motif underlying SATB2 (Special AT-rich Sequence-Binding protein 2) binding showed the top motif to be AT-rich (Supplemental Figure 7B) as expected (Hassan et al., 2010; Naik & Galande, 2019; Savarese et al., 2009; L. Q. Zhou et al., 2012). GREAT analysis of GO-terms associated with ChIP-seq peaks, showed SATB2 binding to be enriched at loci associated with NC development and migration, and Epithelial-to-Mesenchymal Transition (EMT) (Figure 3A). Indeed, ~38.8% (127/327) of known NC-associated genes (Tan et al., 2016), including *sox10, snai2, pax7, chd7, and semaforin (sema3fa*) are bound by SATB2 (Supplementary Table 2). We next performed ChIP-seq for H3K27Ac and H3K9me3 histone marks, in MCR:SATB2 and MCR:EGFP control tumors to investigate the effect of SATB2 overexpression on the chromatin state of SATB2-bound targets. Globally, genome wide GO term analysis of loci with increased H3K27Ac deposition in MCR:SATB2 vs. MCR:EGFP also showed an increased activation of neural crest development (Supplementary Figure 7C). Furthermore, HOMER motif analysis of known motifs shows SATB2 ChIP-seq to be enriched for TFAP2A and RXR motifs, which are both transcription factors involved in neural crest specification (Supplementary Figure 7D). Locally, H3K27Ac deposition in MCR:SATB2 vs. MCR:EGFP around transcriptional start sites (TSS) of SATB2-bound genes suggested an increased chromatin activation state of SATB2-bound NC targets (Figure 3B).

**Figure 3:**
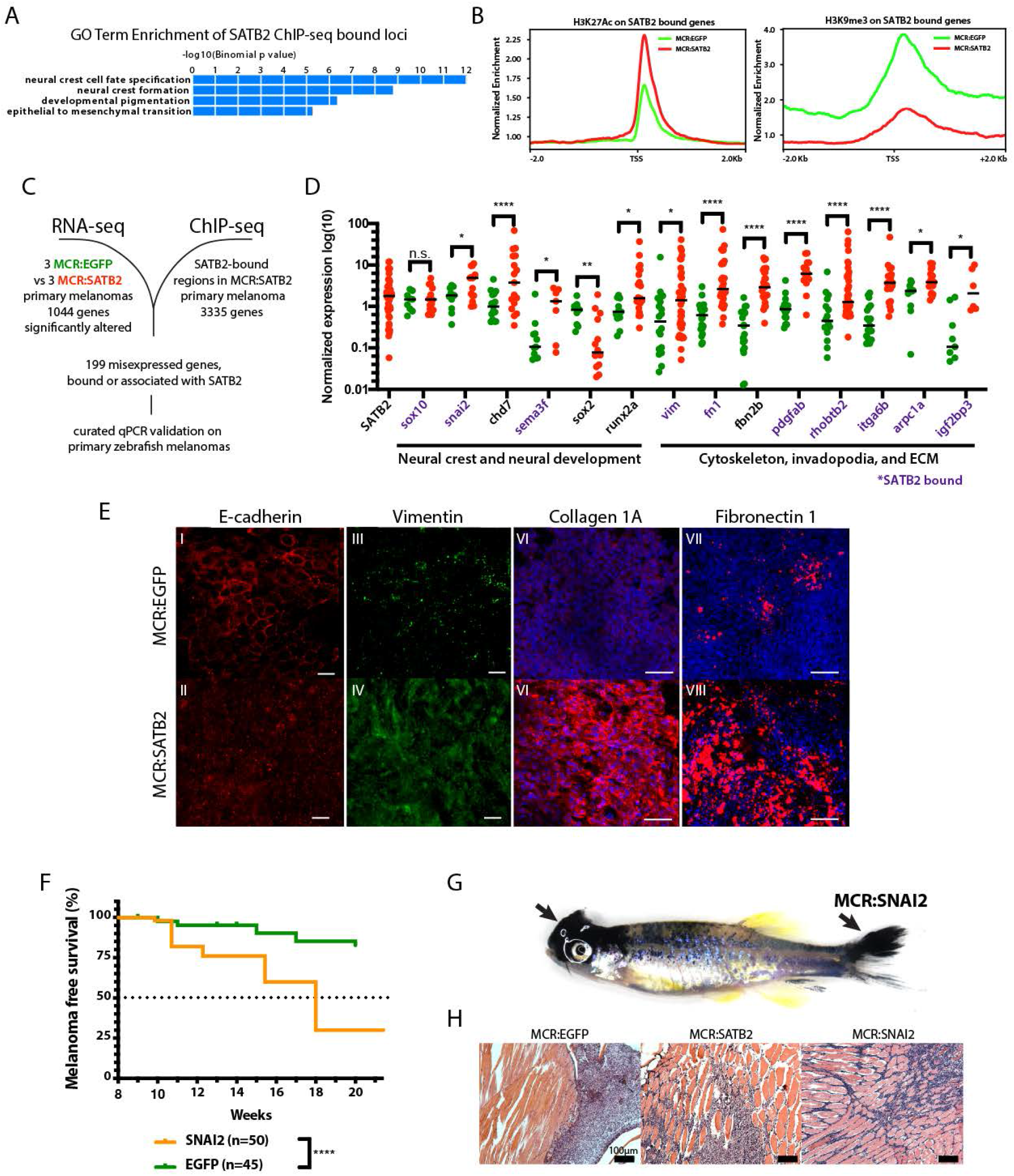
SATB2 binds and regulates EMT and neural crest-associated genes. (A) GO-term enrichment of SATB2-bound loci (GREAT analysis) from anti-SATB2 ChIP-seq on MCR:SATB2 tumors. Genes are defined as bound if SATB2 peaks are found within 3kb of the transcriptional start (TSS) or end site (TES), and the gene body (GB). (B) H3K27Ac and H3K9me3 TSS +/− 2Kb profile of H3K27Ac at SATB2 bound targets comparing MCR:SATB2 in red and MCR:EGFP in green. (C) ChIP-seq and RNA-seq on primary zebrafish tumors was performed to identify SATB2-bound genes that are significantly differentially expressed (P<0.05, q<0.05, FC>|1.5|). (D) qRT-PCR validation of transcriptionally altered SATB2-bound and associated targets associated to neural crest development, and the actin cytoskeleton and extracellular matrix. Symbols indicate single primary tumors, normalized to β-actin. \**P*<0.05, \*\*\**P*<0.0001, \*\*\*\**P*<0.0001. SATB2-bound genes are highlighted in purple. (E) Whole mount primary tumor immunohistochemistry for E-cadherin (red), scale bar is 100 μm, Vimentin (green), scale bar is 100 μm, Fibronectin 1 (red), scale bar is 50 μm, and Collagen 1 (red), scale bar is 50 μm, Dapi (blue). (F) Kaplan-Meier melanoma-free survival curves of MCR:EGFP and MCR:SNAI2 overexpressed in the zebrafish MCR melanoma model (logrank, *p<0.0001****.* (G) Gross anatomy of zebrafish injected with MCR:SNAI2. Arrows (black) show melanomas. (H) Representative H&E histology of MCR:EGFP, MCR:SATB2, and MCR:SNAI2 tumors. Scale bar (black) is 100μm.

SATB2’s binding pattern is consistent with its role as a necessary specifier of cranial migratory neural crest differentiation in exo-mesenchymal tissues in the pharyngeal arches (which will develop into the jaw and teeth), and the consequent cleft palate defects observed in patients with mutated *SATB2* in the human SATB2 syndrome (Zarate & Fish, 2017). Consistent with these prior literature findings we find *satb2* to be highly expressed in a previously published dataset of migrating *crestin:EGFP^+^-NC* cells sorted from zebrafish embryos at the 15 somite stage (Kaufman et al., 2016), when EMT-mediated delamination initiates migration of the neural crest, including cranial (Supplementary Figure 7E). Furthermore, knock-down of *satb2* during embryonic development via microinjection of a validated splicing morpholino (Ahn et al., 2010) in the transgenic neural crest reporter zebrafish line *Tg(sox10:mCherry)* showed a reduction of *sox10* reporter expression and craniofacial abnormalities, but did not affect melanocyte development (Supplementary Figure 7F).

To define the transcriptional effect of SATB2-overexpression we performed RNA-seq on 3 MCR:SATB2 and 3 MCR:EGFP tumors, and correlated SATB2-bound loci with their transcriptional changes in MCR tumors to identify SATB2 target genes that might underlie SATB2’s phenotype (Figure 3C). Given the inter-tumor transcriptional variability of primary zebrafish tumors (evident by qPCR on SATB2 itself in MCR:SATB2 tumors, Figure 3D), we conducted extended validation on a subset of these bound and transcriptionally altered genes, plus some additional manually curated SATB2-bound genes by qPCR on a large set of primary tumors (Figure 3D). Genes that are SATB2-bound [within 3kb of the transcriptional start or end site, and the gene body] or SATB2-associated (predicted nearest gene association), which were altered by RNA-seq revealed that NC, cytoskeleton and extracellular matrix-associated gene programs are highly stimulated by SATB2-overexpression (Figure 3D, Supplementary Table 2). *chd7* and *snai2* in particular have been described to be key regulators of the migratory NC (Bajpai et al., 2010; Okuno et al., 2017; Schulz et al., 2014), while *pdgfab* has been implicated in podosome and invadopodia formation (Ekpe-Adewuyi, Lopez-Campistrous, Tang, Brindley, & McMullen, 2016; Murphy & Courtneidge, 2011; Murphy et al., 2011; Paz, Pathak, & Yang, 2014). SATB2-binding to targets such as *pdgfab* (Supplemental Figure 7G), results in increased H3K27Ac deposition along the promoter, TSS region and gene body in MCR:SATB2 compared to MCR:EGFP, respectively.

To investigate whether in MCR:SATB2 tumors cell fate rewiring functionally induces an EMT-like phenotype-switch reminiscent of the cranial neural crest development, we performed whole mount immunofluorescence for EMT markers on primary zebrafish melanomas. Indeed, compared to MCR:EGFP, MCR:SATB2 tumors showed mislocalized E-cadherin, and an increased protein expression of Vimentin and extracellular matrix attachment proteins Fibronectin 1 *(fn1a/b)* and Collagen 1 *(col1a1a/b)* (Figure 3E). Given the known role of *snai* family of proteins in regulating EMT we tested whether SATB2’s direct downstream target *snai2* could recapitulate the MCR:SATB2 phenotype *in vivo*. Overexpression of SNAI2 resulted in a significant acceleration of tumor onset compared to the EGFP negative control (Figure 5F), albeit milder than SATB2 (MCR:SNAI2 median onset 18 weeks vs 12 weeks for MCR:SATB2). Similar to MCR:SATB2 (Figure 1D), and unlike MCR:EGFP, MCR:SNAI2 fish showed development of multiple tumors (Figure 5G), with a histopathologically invasive morphology (Figure 5H). Overall, SATB2 binds and activates chromatin at a subset of neural crest-related loci including *snai2*, resulting in an EMT-like phenotype. Overexpression of the human ortholog of downstream target *snai2* partially recapitulates SATB2’s phenotype in the MCR model.

### SATB2-induced program is conserved across zebrafish and human melanoma and overlaps with known drug resistance transcriptional states

To ascertain whether SATB2-induced transcriptional changes were conserved across species, we inducibly overexpressed SATB2 in primary human melanocytes and human melanoma cell lines, and performed qPCR for selected orthologues of genes validated in zebrafish. qPCR analysis of SATB2 target genes after 48 hours of culture in the presence of doxycycline on iSATB2 melanoma cell lines (SKMEL2, A375 and SKMEL28 iSATB2), and iSATB2 untransformed primary human melanocytes (HEMA-LP) showed a similar increase in SNAI transcription factors (Figure 4A). Cell line specific differences in overall *SNAI1* or *SNAI2* (both *snai2* orthologs) induction levels may reflect context dependence (e.g. oncogene or cell line baseline status) (Nieto, Huang, Jackson, & Thiery, 2016). To analyze global transcriptional changes induced by SATB2 in human melanoma, we selected SKMEL2 iSATB2, since this cell line had the lowest endogenous SATB2 level and showed the strongest invadopodia induction (Figure 2D-E and Supplementary Figure 4B). We conducted RNA-seq on SKMEL2 iSATB2 after 48 hours of treatment with doxycycline. Both Gene Set Enrichment Analysis (GSEA) Hallmark pathways analysis and Ingenuity Pathway Analysis (IPA) (Figure 4B) of significant genes, showed similar effects of SATB2 overexpression between human melanoma cell line (SKMEL2 iSATB2 *p*<0.01; *q*<0.01; FC>2) and zebrafish tumors (MCR:SATB2 vs. EGFP *p*<0.01; *q*<0.01; FC>1.5) with strong similarity in the top significant pathways across human and zebrafish melanoma. These overlaps included pathways related to EMT/mesenchymal activation, osteoblast changes, axonal guidance, semaphorins, cancer metastasis, WNT, and TGFB signaling (Figure 4B). SATB2-overexpression (SKMEL2 iSATB2 RNA-seq) induced consistent transcriptional changes in several NC and invadopodia regulators, including activation of human orthologs of *pdgfab* and *snai2* (Figure 4C). These similarities across species, overexpression systems, and *in vitro* vs. *in vivo* conditions consolidate our findings.

**Figure 4:**
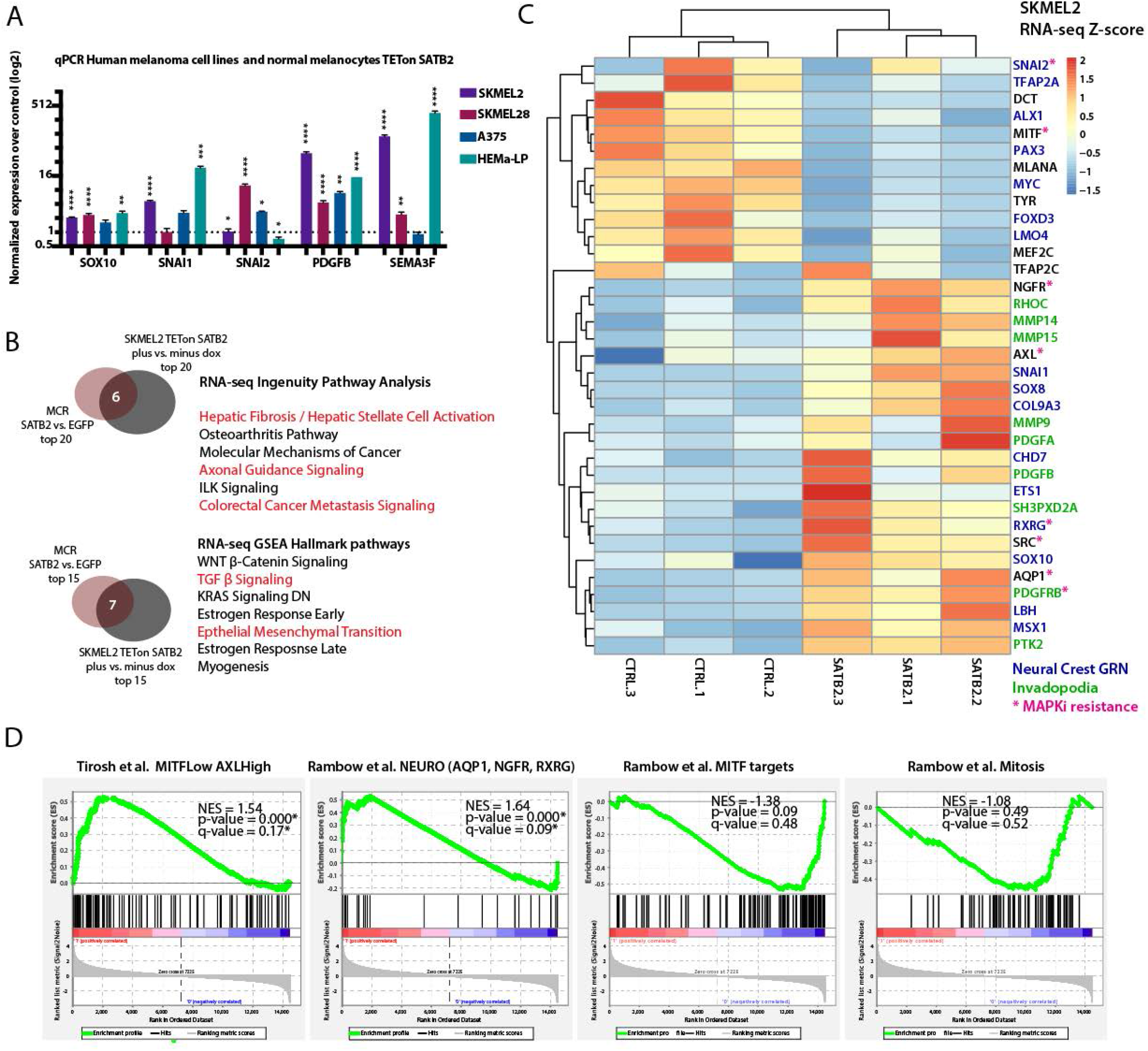
Cross-species conservation of SATB2-overexpression induced transcriptional program and known drug resistant cell states. (A) Overexpression of SATB2 with TETon (iSATB2) in normal human epidermal melanocytes (HEMA-LP) and in SKMEL2, SKMEL28 and A375 melanoma cells is sufficient to induce transcriptional activation of *SOX10, SNAI1/2, PDGFB,* and SEMA3F mRNA. qRT-PCR data was normalized to *beta-actin,* and fold change was determined compared to un-induced controls. \**P*<0.05, \*\**P*<0.001, \*\*\**P*<0.0001, \*\*\*\**P*<0.0001. (B) Conservation of top regulatory pathways identified by RNA-seq Ingenuity Pathway Analysis and Gene Set Enrichment Analysis (Hallmark pathways) comparing MCR:SATB2 vs MCR:EGFP zebrafish tumors with iSATB2 (plus DOX vs. minus DOX) SKMEL2 human melanoma cell line. For IPA humanized significant zebrafish genes were used (P<0.05, q<0.05, FC>|1.5|=1044 zf-> 840 unique human orthologs), and significant SKMEL2 iSATB2 genes (P<0.05, q<0.05, FC>|1|). (C) Curated heatmap of altered genes in RNA-seq of human SKMEL2 cells overexpressing SATB2 (plus DOX vs. minus DOX FC>1; p<0.01) that have been involved in neural crest, Rambow Neural crest-like state (Rambow et al., 2018), PDGF-PDGFR-SRC invadopodia cascade, AXL/MITF state (Tirosh et al., 2016) and melanocyte differentiation. All genes shown are significantly altered except for TFAP2C. Normalized RNA-seq heatmap of expression fold-change Z-score is plotted. Genes part of the neural crest gene regulatory network (GRN) are highlighted in blue, invadopodia related genes are highlighted in green, and MAPK inhibitor resistance genes are highlighted in magenta. (D) Gene set enrichment analysis (GSEA) of 1000 MSigDB signatures including the states described by Tirosh *et a/*.(Tirosh et al., 2016) and Rambow *et al(Rambow et al., 2018).*

To assess how the SATB2 program relates to previously described transcriptional states in melanoma we conducted GSEA analysis in SKMEL2 iSATB2 for known signatures: (1) a mesenchymal signature (Verfaillie et al., 2015), (2) MITF^low^/AXL^high^ drug resistance state (Tirosh et al., 2016), and (3) neural crest state and other transcriptional signatures in different tumor cell subpopulations (Rambow et al., 2018^)^. This analysis showed a significant overlap of the SATB2-induced transcriptional program with the less differentiated MITF^low^/AXL^high^ state (NES=1.54, *p*=0.000, *q*=0.17), and the Rambow neural crest-like/minimal residual disease MITF^low^/NGFR1^high^/AQP1^high^ state driven by *RXRG* (NES=1.64, *p*=0.000, *q*=0.09) (Figure 4D), which have both been previously described in metastatic human melanoma to correlate with a self-renewing MAPK inhibitor drug resistant state (Rambow et al., 2018; Tirosh et al., 2016). The key marker genes of these states (i.e. *AQP1, RXRG, NGFR, AXL,* Figure 4C) and additional genes involved in MAPK resistance (i.e. SRC and PDGF pathway members (Ekpe-Adewuyi et al., 2016; Nazarian et al., 2010; Rebecca et al., 2014), Figure 4C) and RXR related pathways (Supplementary Figure 8A-B) are significantly induced by SATB2 overexpression. Overall, SATB2 drives a transcriptional induction of invadopodia, EMT and neural crest regulators in zebrafish and human melanoma alike, consistent with the transcriptional changes and phenotypes described above and its known role in development. Furthermore, the SATB2-induced transcriptional program shows strong overlaps with known drug resistance transcriptional states in melanoma.

### SATB2 increases tumor propagating potential and resistance to MAPK inhibition in vivo

Given the overlap between the SATB2-induced program and the known minimal residual disease and MAPK inhibition resistant states, we asked whether SATB2 functionally confers enhanced self-renewal or resistance to MAPK inhibition. To address the former, we performed a limiting dilution tumor propagation assay injecting 300,000, 3,000, 300 or 30 primary pigmented melanoma cells from MCR:EGFP (n=4) and MCR:SATB2 (n=5) donor tumors, into the dorsum of the zebrafish (Figure 5A). While MCR:EGFP did not display engraftment with fewer than 3000 cells (0/28), 44% (4/9) of MCR:SATB2 engrafted with 30 cells (Figure 5B). SATB2 donors have a significantly higher estimated fraction of tumor propagating cells (median estimate 1/21.6 cells for MCR:SATB2 vs. 1/10,879 cells for MCR:EGFP, Mann Whitney *p*=0.0159*) (Figure 5C).

**Figure 5:**
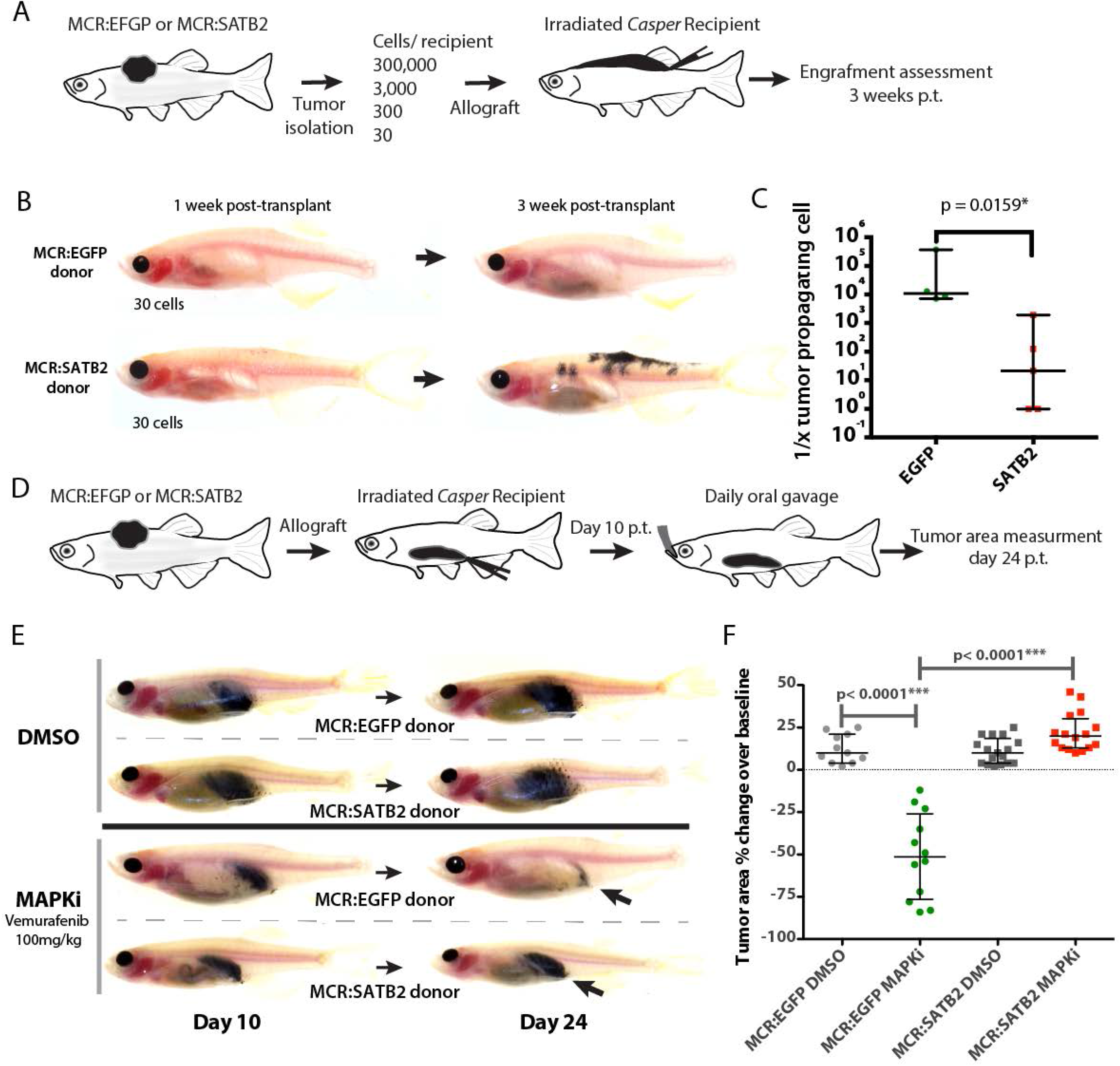
MCR:SATB2 tumor allografts have higher tumor propagating potential and resistance to Vemurafenib *in vivo*. A) Schematic overview of the subcutaneous primary melanoma limiting dilution transplantation model in transparent *casper* zebrafish. 300,000, 3,000, 300 or 30 MCR:SATB2 or MCR:EGFP primary pigmented primary melanoma cells were transplanted into a subcutaneous space in running along the dorsum of irradiated *casper* recipients, and assessed for engraftment 1, 2 and 3 weeks post-transplant. (B) Representative images of MCR:EGFP and MCR:SATB2 recipients transplanted with 30 cells. (D) Engraftment of surviving recipients at 3 weeks post-transplant was used to estimate the frequency of tumor propagating cell potential with Extreme Limiting Dilution Analysis (ELDA). Estimated frequency of tumor propagating cells between MCR:EGFP and MCR:SATB2 donors (Mann-Whitney, *p*<0.0159*). (D) Schematic overview of *in vivo* drug treatment of allotransplanted primary MCR:EGFP or MCR:SATB2 tumors with Vemurafenib. Recipients *casper* fish were treated by daily oral gavage with 100mg/kg of drug or vehicle control from day 10 post-transplant to day 24 post-transplant. Animals were imaged at day 10 and 24 and the pigmented tumor area was estimated using digital calipers to assess treatment response. (E) Representative image of individual DMSO or Vemurafenib treated animals transplanted with MCR:SATB2 or MCR:EGFP cells at day 10 and day 24. (F) Mean normalized area percent change in the treated cohorts: MCR:EGFP DMSO (n=11) +12.45%, SEM 2.55% vs. MCR:EGFP MAPKi (n=12) −50.67%, SEM 7.26% vs. MCR:SATB2 DMSO (n=17) +11.06%, SEM 1.84% vs. MCR:SATB2 MAPKi (n=16) +22.06%, SEM 2.81%. Unpaired 2-tailed t-test *p* values of pairwise comparisons are shown.

To test whether the SATB2-induced program confers resistance to MAPK inhibition, we utilized an established *in vivo* drug treatment assay(Dang, Henderson, Garraway, & Zon, 2016) where primary zebrafish melanoma are allotransplanted in *casper* recipients and administered a MAPKi (Vemurafenib 100mg/kg) daily via oral gavage (Figure 5D). Analysis of tumor growth by measuring tumor area with digital calipers after two weeks (day 24 post-transplant) of treatment with Vemurafenib or a DMSO vehicle control showed a complete lack of response in MCR:SATB2 compared to Vemurafenib sensitive MCR:EGFP tumors (Figure 5E-F) (MCR:EGFP MAPKi vs. MCR:SATB2 MAPKi p< 0.0001****). Our data functionally validate SATB2 as a driver of enhanced tumor propagation and drug resistance *in vivo.*

## Discussion

In this study we investigated the effect of epigenetic and transcriptional regulators on melanoma development, by *in vivo* perturbation in endogenous tumors. We screened 80 human chromatin factors using a pooled screening approach in a transgenic zebrafish melanoma model driven by the most common melanoma oncogene, *BRAF^V600E^*, and loss of the tumor suppressor *tp53* (Cancer Genome Atlas, 2015). Through single factor validation, we identified transcriptional regulator SATB2 as a potent accelerator of melanoma onset in zebrafish (Figure 1B-C, Supplementary Video 1). MCR:SATB2 transgenic animals, in addition to developing tumors faster, had a high prevalence of tumor burden in internal organs, which is uncommon in MCR:EGFP controls, even in animals with very large tumors (Figure 1D-G, Supplementary Figure 1D-E). This has not been previously observed with several other accelerators or drug resistance drivers that we and others have identified using the MiniCoopR model (Ablain et al., 2018; Ceol et al., 2011; Dang et al., 2016; Fazio et al., 2020; Iwanaga et al., 2020; Kaufman et al., 2016; Venkatesan et al., 2018).

This phenotype could be explained by a proliferation effect resulting in multiple parallel tumor initiation events in both dermal and internal melanocytes (Kapp et al., 2018), or by an early invasive migration and metastasis phenotype. We did not observe proliferation differences between MCR:SATB2 and MCR:EGFP (Supplementary Figure 3A), nor in a panel of three human melanoma cell lines upon inducible overexpression of SATB2 (Supplementary Figure 3B-C). On the other hand SATB2 overexpression in cultured zebrafish tumors (Figure 2A and Supplementary Figure 4A-D) and human melanoma cell lines (Figure 2A, Supplementary Figure 4A-D, and Supplementary Video 2) resulted in increased invasion *in vitro*, and MCR:SATB2 zebrafish primary tumor allografts in irradiated optically clear *casper* recipients had a much higher distant and internal metastasis rate compared to MCR:EGFP controls (Figure 2F-J, Supplementary Figure 5D, and Supplementary Video 3-4), suggesting the latter hypothesis to be more likely at the basis of our observations in tumor-bearing MCR:SATB2 transgenic fish.

Our chromatin and transcriptional analysis via ChIP-seq, and RNA-seq followed by qPCR validation on MCR:SATB2 vs. MCR:EGFP control tumors showed SATB2 to directly bind and transcriptionally induce several genes related to neural crest development (*snai2* and *chd7*) (Bajpai et al., 2010; Betancur, Bronner-Fraser, & Sauka-Spengler, 2010; Mayor & Theveneau, 2013; Schulz et al., 2014), EMT (*vim* and *fn1*) (Betancur et al., 2010), and invadopodia formation *(pdgfab* and *rhobtb2)* (Maheswaran & Haber, 2015; Mayor & Theveneau, 2013; Murphy & Courtneidge, 2011; Nieto et al., 2016; Paz et al., 2014). Functionally, an EMT-like phenotype and ECM remodeling in MCR:SATB2 vs. MCR:EGFP tumors was also apparent at the protein level (Figure 3E), and SATB2 overexpression induced the formation of functional invadopodia with ECM degrading capacity in zebrafish tumors, primary cell lines, and human melanoma cell lines alike (Figure 2A-E, Supplementary Figure 4B-E, and Supplementary Video 2). This neural crest mesenchyme-like program outlined above is consistent with SATB2’s known roles in the development of exo-mesenchymal derivatives of the cranial neural crest (Supplementary Figure 7E-F) (Ahn et al., 2010; Hassan et al., 2010; Kikuiri et al., 2018; Nanni et al., 2019; Rainger et al., 2014; Sheehan-Rooney et al., 2010; Sheehan-Rooney et al., 2013; Zarate et al., 2019; Zarate & Fish, 2017; P. Zhou et al., 2016), neuronal development and axon guidance (Gyorgy, Szemes, de Juan Romero, Tarabykin, & Agoston, 2008; McKenna et al., 2015; Shinmyo & Kawasaki, 2017), and regulation of EMT/invadopodia in other solid tumors (Gan et al., 2017; Mansour et al., 2015; Naik & Galande, 2019; Wu et al., 2016; Xu et al., 2017; Yu, Ma, Ochoa, et al., 2017). EMT and invadopodia formation are, in fact, inherent to migration of the neural crest itself, since neural crest cells are specified in proximity of the neural plate, undergo EMT and rely upon podosomes, termed invadopodia in cancer cells, to remodel the ECM and migrate through the body (Bailey, Morrison, & Kulesa, 2012; Betancur et al., 2010; Mayor & Theveneau, 2013; Murphy & Courtneidge, 2011; Murphy et al., 2011). Reactivation of neural crest regulator *snai2* has been shown to increase the transformation and migration potential of melanoma (Gupta et al., 2005; Shirley et al., 2012) Overexpression of human SNAI2 in the MiniCoopR model similarly resulted in faster tumor development compared to EGFP controls (*P*<0.0001), although not to the same extent as SATB2 (MCR:SNAI2 median onset 18 weeks, MCR:SATB2 median onset 12 weeks, MCR:EGFP median onset ~22 weeks in both experiments), (Fig 1C and Fig 3F-G) and similar invasive histological features compared to MCR:SATB2 tumors (Figure 3H). Taken together, these data suggest that downstream SATB2 target *snai2* is partially responsible for the SATB2 phenotype, but that other genes in the program contribute to the full phenotype spectrum.

To further assess the conservation and relevance of our findings to human melanoma, we conducted qPCR analysis on a panel of iSATB2 human melanoma cell lines and primary cultured melanocytes (HEMa-LP). qPCR of the human orthologs of *snai2, pdgfab,* and *sema3f* overall showed a similar induction as zebrafish tumors, with some variation across cell lines (Figure 3D and 4A). For an unbiased global transcriptional comparison, we conducted RNA-seq on human melanoma cell line SKMEL2 iSATB2 and identified transcriptional differences induced by SATB2 overexpression (with/without Dox induction for 48 hours). Significant differentially regulated genes from SKMEL2 iSATB2 were compared with zebrafish primary tumor RNA-seq (MCR:SATB2 vs. MCR:EGFP) using both Gene Set Enrichment Analysis (GSEA) and Ingenuity Pathway Analysis (IPA). Despite 1) the ortholog prediction constraints between human and zebrafish datasets that may have resulted in some expected loss of information, 2) the variation due to species genetic and functional conservation differences, and 3) the *in vivo* vs. *in vitro* conditions, we see strong conservation across the two models in the top altered pathways (Figure 4B). Overall, our study shows a robust transcriptional and phenotypic conservation of SATB2 overexpression effects in melanoma across human and zebrafish (Figure 2A-E, Supplementary Figure 4B-E, Figure 4B-C, Supplementary Figure 8A-B, and Supplementary Figure 3A,C). The SATB2-driven program we describe here (Figure 4C-D) significantly overlaps with the recently reported MITF^low^/AXL ^high^ state (Tirosh et al., 2016), and the neural crest-like MITF^low^/NGFR1^high^/AQP1^high^ MAPK inhibitor resistance less proliferative states (Rambow et al., 2018). Indeed, SATB2 overexpression in SKMEL2 (Figure 4C, Supplementary Figure 8A) induced up-regulation of AXL, AQP1, RXRG, NGFR, PDGFRB, and SRC, which have all been reported to drive resistance to MAPK inhibitors (Boshuizen et al., 2020) (Nazarian et al., 2010) (Fallahi-Sichani et al., 2017). Interestingly, SATB2 has been reported as a primary hit in an unbiased overexpression screen for MAPK inhibitor resistance drivers in human melanoma cell lines *in vitro* (Johannessen et al., 2013). We sought out to functionally validate SATB2 as a resistance driver by conducting *in vivo* limiting dilution transplants and drug treatments with Vemurafenib using established assays in zebrafish allografts (Dang et al., 2016; Heilmann et al., 2015) and showed MCR:SATB2 tumors to have increased tumor propagating potential (Figure 5A-C), and primary resistance to MAPK inhibitors *in vivo* (Figure 5D-F).

Analysis of publicly available genomic datasets showed SATB2 to not be recurrently altered in the TCGA dataset (Consortium, 2020), but its amplifications is observed in ~4-8% of patients in 3 independent datasets of metastatic melanoma (Supplementary Figure 6C) (Hugo et al., 2015; Snyder et al., 2014; Van Allen et al., 2014), with a similar fraction of SATB2 high-expressing patients having a higher risk of metastasis-related poor outcome in two additional datasets (Supplementary Figure 6D).

In summary, our work identifies SATB2 as a driver of invasion and resistance to Vemurafenib treatment in melanoma. Yet, further studies will be needed to assess the prevalence of SATB2 expression changes in targeted therapy and immunotherapy clinical settings. Furthermore, we show in an endogenous transgenic tumor model that SATB2 transcriptional rewiring of melanoma towards a state similar to the less proliferative neural crest-like state described by Rambow and colleagues (Rambow et al., 2018) can drive accelerated tumor development and invasion. Our work reinforces the idea that melanoma phenotype switching and transcriptional plasticity might dynamically select a dominant state depending on environmental constraints on proliferation during different stages of the tumor’s natural history (Ahmed & Haass, 2018; Boumahdi & de Sauvage, 2020; F. Z. Li, Dhillon, Anderson, McArthur, & Ferrao, 2015; Marine, Dawson, & Dawson, 2020).

## Acknowledgments

We thank Serine Avagyan for helpful discussion, Elliott Hagedorn for helpful discussion and time-lapse imaging support, Cristine Lian for histopathology discussion, and Rachel Fogely for ChIP-seq guidance. EvR was supported by a Netherlands Organization for Scientific Research (NWO) Rubicon fellowship, and a Dutch Cancer Foundation (KWF) fellowship for Fundamental Cancer Research. MF was supported by Boehringer Ingelheim Fonds. LIZ is supported by: R01 CA103846, MRA (Zon, Garraway), The Starr Cancer Consortium and the Ellison Foundation.

## Author Contributions

MF designed the study, conducted and interpreted experiments, and wrote the manuscript. EvR designed the study, conducted and interpreted experiments and wrote the manuscript. GvdH contributed to the chromatin factor screen and performed experiments. AT, JM and RM contributed to animal husbandry, cell culture maintenance and transgenic injections. MD performed subcutaneous zebrafish allograft and drug treatment assays. JM performed histopathological analysis of zebrafish melanoma. JA designed and generated CRISPR constructs. PP and CK provided experimental support. RW assisted with RNA-seq analysis and developed the zebrafish melanoma cell culture conditions and allotransplantation in *casper.* SY and YZ performed bioinformatical analysis on the RNA-seq and ChIP-Seq data. LIZ conceived the study and interpreted experiments.

## Methods

### Zebrafish melanoma model and MiniCoopR system

Experiments were performed as outlined by Ceol *et al.* (Ceol et al., 2011). MiniCoopR (MCR) expression constructs were created by MultiSite Gateway recombination (Invitrogen) using fulllength human open reading frames. Briefly, pools of five similarly sized MiniCoopR constructs (5 pg each), or single factors (25 pg) including MCR alone, MCR:EGFP *(Ceol et al., 2011),* MCR:SATB2 (IOH46688, Invitrogen), MCR:SNAI2 (pCMV-SPORT6-SNAI2 [HsCD00327569]),and MCR:SETDB1 *(Ceol et al., 2011),* were microinjected together with 25 pg of Tol2 transposase mRNA into one-cell *Tg(BRAF^V600E^);p53^-/-^; mitf^-/-^* zebrafish embryos. Embryos were scored for melanocyte rescue at 48-72 hours post-fertilization, and equal numbers were raised to adulthood (15-20 zebrafish per tank), and scored weekly (from 8-12 weeks post-fertilization) or bi-weekly (> 12 weeks post-fertilization) for the emergence of raised melanoma lesions. The minimum cohort size per injection was 40 animals as previously defined (Ceol et al., 2011). We pre-established criteria to exclude/censor animals if they died without having been examined for tumor formation. No randomization or blinding was used. Kaplan-Meier survival curves were generated in GraphPad Prism, and statistical difference was determined by a log-rank (Mantel-Cox) test. For subsequent expression, binding and histopathological analyses, large tumors were isolated from MCR:SATB2 (9-18 weeks post-fertilization), MCR/MCR:EGFP (14-28 weeks postfertilization), due to later tumor onset and slower progression compared to MCR:SATB2 and MCR:SNAI2 (14-20 weeks post-fertilization). Zebrafish were maintained under IACUC-approved conditions.

### CRISPR/Cas9 inactivation of *satb2*

To specifically inactivate *satb2* in melanocytes, we engineered the MiniCoopR vector to express Cas9 under the control of the melanocyte-specific *mitfa* promoter, and a gRNA efficiently mutating *satb2* off a *U6* promoter (Ablain et al., 2015). *Cas9* mRNA was produced by *in vitro* transcription from a pCS2 *Cas9* vector (Jao et al., 2013) using mMESSAGE mMACHINE SP6 kit (Invitrogen). The *satb2* gRNA 5’-GGATGGGCAGGGGGTTCCAG-3’ was generated following established methods (Gagnon et al., 2014). To validate targeting, 600 pg of *Cas9* mRNA and 25 pg of gRNA were injected into embryos of the AB strain, and the T7E1 assay was performed as reported (Kim et al., 2009), using the following primers: 5’-CCTACCTCAATCCACTCTTT-3’ and 5’-GCTGCACCAAGAAACTACAA-3’. Verified gRNAs against *satb2,* and a *p53* control (Ablain et al., 2015) were injected into one-cell stage *Tg(mitfa:BRAF^V600E^); p53^-/-^; mitfa^-/-^* embryos, and tumor formation was monitored.

### Morpholino knockdown *of satb2*

To address whether SATB2 is necessary for neural crest and melanocyte development, 4 pg of a splicing morpholino against *satb2* (GCAGTGTTGAACTCACCATGAGCCT, Ahn et al., 2010) was injected into one-cell stage *TG(sox10:mCherry)* embryos. At 3 dpf, injected and uninjected control embryos were scored for melanocyte and cranio-facial abnormalities using light and fluorescence microscopy. The experiment was repeated 3 times, with an average clutch size of 40 embryos per experiment.

### Zebrafish primary tumor and cell line transplants

Pigmented primary melanomas were excised from euthanized MCR/MCR:EGFP and MCR:SATB2 zebrafish, and dissociated in 50% Ham’s-12/DMEM medium containing 0.075 mg/mL liberase (Promega) for 30 minutes at RT, with periodical manual disaggregation using a razor blade. The dissociation medium was inactivated by the addition of 3 x 5 ml 50% Ham’s-12/DMEM with 15% heat-inactivated Fetal Bovine Serum. To obtain a single cell suspension, cells were passaged through a 40 μm filter into a 50 mL falcon tube, and pelleted by centrifugation at 500 x g for 5 minutes. Cell numbers were determined, and a cell suspension of 100,000 cells/μl in PBS was made, and kept on ice. *casper* recipients received a split-dose irradiation of 15 Gy over two consecutive days prior to transplantation (Heilmann et al., 2015). During transplantation, irradiated *casper* recipients were anesthetized in 0.4% MS222, mounted in a moist sponge. Using a Hamilton syringe, 300,000 pigmented primary tumor cells, or 250,000 MCR:EGFP (zmel1) or MCR:SATB2 (45-3) cells were injected into a confined interstitial subcutaneous space that runs along the dorsum of the zebrafish. Five to seven primary tumors were used per cohort, with each individual tumor being transplanted into 6-10 *casper* recipients. Metastatic progression, as defined by the formation of distant metastasis that have spread passed the anatomical midline, was monitored weekly or at the experimental end point at 3-3.5 weeks post-transplant, and photographically recorded using a Nikon D3100 DSLR camera. Limiting dilution experiments were performed in a similar fashion by using multiple dilutions from each individual donor and transplanting cohorts of 10 recipients per each experiment arm. Overall 4 MCR: EGFP donors and 5 MCR: SATB2 donors were transplanted in the course of 3 independent experiments. Recipients were monitored weekly for engraftment or at the experimental end point at 3 weeks post-transplant, and photographically recorded using a Nikon D3100 DSLR camera. The percentage of engraftment and *n* was used as input to estimate the fraction of tumor propagating cells using Extreme limiting dilution analysis ELDA (Hu and Smyth, 2009) Statistical difference across groups was calculated with a Mann-Whitney test using GraphPad Prism. No blinding was performed, and the group size was not predetermined statistically. Statistical significance for all above experiments was determined by an unpaired two-tailed *t*-test unless otherwise specified, using GraphPad Prism. Drug treatment with Vemurafenib (100mg/Kg dissolved in DMSO) or with equal amount of DMSO dissolved in water was performed via daily oral gavage for 14 days starting on day 10 post-transplant on irradiated *casper* recipients allografted in the peritoneum with 500,000 pigmented primary tumor cells. Recipients were monitored at day 10 post transplant before starting treatment, and at the experimental end point at day 24 post-transplant (day 14 of treatment), and photographically recorded using a Nikon D3100 DSLR camera. Tumor response was assessed as previously described (Dang et al., 2016). Tumor area was measured at day 10 and day 24 using a traceable digital caliper (FisherScientific, #14-648-17). The pigmented tumor area was calculated by the longest measured length and width of the tumor. The drug response was quantified via the change from baseline tumor area using the RECIST (Response Efficacy Criteria in Solid Tumors). The response rate for experimental cohorts was depicted via waterfall plots, and *t*-test statistics were applied for significance as previously described (Dang et al., 2016).

### Whole-mount tumor immunohistochemistry

Zebrafish where euthanized on ice, melanomas were surgically removed, and fixed in either 4% paraformaldehyde (PFA) or DENTS (80% methanol, 20% DMSO) overnight at 4°C. Tumors were washed at least 3 times with PBS, and mounted in 4% agarose in PBS for Vibratome sectioning. 100-150 μM sections were cut using the Microm HM650V Vibratome. Sections were next processed for immunohistochemistry. PFA fixed samples were permeabilized by digestion with 15 ug/ml Proteinase K in PBS-0.1% Tween for 30 minutes at 37°C, or 0.4% Triton in PBS for 10 minutes at room temperature. After washing 3 times with PBS-0.1% Tween for 10 minutes, samples were blocked (10% lamb serum, 1% DMSO in PBS-0.1% Tween) for at least one hour, and incubated with primary antibodies overnight at 4°C. Antibodies used were: anti-E Cadherin (1:100, DENTs fixation; ab11512, Abcam), Anti-Vimentin (1:200; sc-6260, Santa-Cruz), antiCollagen I (1:250; ab23730, Abcam), anti-Fibronectin 1 (1:250; F3648, Sigma Aldrich) and anti-Cortactin (p80/85), clone 4F11 (1:100; 05-180, EMD Millipore). After extensive washes with PBS-0.1% Tween, and blocking for one additional hour, samples were incubated in appropriate secondary Alexa-Fluor antibodies (1:500 Invitrogen) overnight at 4°C. Nuclei were counterstained with 1:1000 DAPI, and sections were mounted on a microscope slides in Slow Fade Gold (Invitrogen). Confocal images were collected on a Nikon C2si Laser Scanning Confocal using 40x water or 63x oil immersion objectives.

### Tumor histology and proliferation index

Zebrafish were fixed in 4% paraformaldehyde, decalcified, paraffin embedded, sectioned (7 μm) and Hematoxylin and Eosin (H&E) stained by the DFCI/HCC Research and BWH pathology cores, using standard procedures. Transmitted light images were collected on a Nikon C2si Laser Scanning Confocal using a Hamamatsu camera. To determine proliferation rates, slides were incubated with Anti-Histone H3 (phospho S10) (1:250; ab5176, Abcam), and colorimetrically stained with DAB following standard procedures. The average number of PH3-positive cells per tumor was determined by counting five high power field images (40-60X) per tumor. An unpaired tailed Student’s T-test was used to determine significance.

### Western Blot

Adherent cells were scraped, and zebrafish tumors were mechanically homogenized in RIPA lysis buffer containing 1:100 protease inhibitors (P8340, Sigma-Aldrich) and 20 μM N-ethylmaleimide (Sigma-Aldrich). Lysates were incubated for 20 minutes on ice, and spun down for 10 minutes at 14,000 RPM at 4°C. Samples were denatured by adding Laemmli sample buffer (BioRad) with 5% β-mercaptoethanol (Sigma-Aldrich), and boiled at 95°C for 5 minutes prior to loading. Protein concentrations were determined using the DC protein assay (BioRad). Proteins (20ug) were separated on a 4-20% mini-PROTEAN TGX (BioRad) precast gel, and transferred onto a nitrocellulose membrane using the iBlotting system (Invitrogen). Primary antibodies used were: Anti-SATB2 (zebrafish tumors, 1ug/ml; Ab51502, Abcam), Anti-SATB2 (human cells, 1:200; HPA029543, Sigma Aldrich) and Anti-Beta Actin (1ug/ml; A2228, Sigma-Aldrich) as a loading control. Additional antibodies used against SATB1 and SATB2 are listed in Supplementary Figure 5A-B. Protein bands were detected by rabbit anti-mouse HRP (1:20,000, Pierce) or swine antirabbit HRP (1:20,000, Pierce).

### RNA extraction, quantitative RT-PCR analysis and RNA-seq sample preparation

Zebrafish tumors were isolated and mechanically homogenized in 350 μl RTL buffer (Qiagen) containing β-mercaptoethanol, on ice for 20 seconds. Adherent cells (zebrafish and human cell culture) were washed twice with ice cold PBS on ice, and cells were scraped in RTL buffer containing β-mercaptoethanol (Sigma-Aldrich). Tumor and cell lysates were next transferred onto a QiaShredder column (Qiagen). RNA isolation was performed using the RNA micro plus kit (Qiagen), according to the manufacturers instruction. RNA quality was determined using a Nanodrop. For RNA-seq on primary zebrafish melanomas, additional quality control of the total RNA was performed on and Fragment Analyzer. Total RNA was depleted of ribosomal RNA with the RiboZero gold kit (Epicentre), and enriched mRNA was applied to library preparation according to manufacturer’s protocol (NEBNext Ultra). After repeated quality control for an average DNA input size of 300 base pairs (bp), samples were sequenced on a HiSeq Illumina sequencer with 2 × 100-bp paired-end reads. For qPCR, cDNA was synthesized with the SuperScript III Kit (Life Technologies) using a 1:1 mixture of random Hexamers and OligoDT. Quantitative PCR was performed on a BioRad iQ5 real-time PCR machine using the Ssofast EvaGreen Supermix (BioRad). The △Ct or △△Ct methods were used for relative quantification. The average of 2 independent experiments is shown for SKMEL2 and a single experiment is shown for primary human melanocytes. qPCR primers were designed using GETprime (Gubelmann et al., 2011) or qPrimerDepot (https://primerdepot.nci.nih.gov). For primer sequences see Supplementary Table 5.

### Zebrafish primary melanoma cell culture

Zebrafish primary melanoma cell lines were generated as described(Heilmann et al., 2015). MCR alone (CK5), MCR:EGFP (zmel1), and MCR:SATB2 (45-3 and 63-4) cell lines were cultured in DMEM medium (Life Technologies) supplemented with 10% heat-inactivated FBS (Atlanta Biologicals), 1X GlutaMAX (Life Technologies) and 1% Penicillin-Streptomycin (Life Technologies), at 28°C, 5% CO_2_. Zebrafish melanoma lines were authenticated by qPCR and Western for human SATB2 or EGFP transgene expression, and periodically checked for mycoplasma using the Universal Mycoplasma Detection Kit (ATCC).

### Human melanoma and primary melanocyte cell culture

Human melanoma cell lines A375, SKMEL2, and SKMEL28 were obtained from the ATCC, and grown DMEM medium (Life Technologies) supplemented with 10% heat-inactivated FBS (Atlanta Biologicals), 1X GlutaMAX (Life Technologies) and 1% Penicillin-Streptomycin (Life Technologies), and grown at 37°C, 5% CO_2_. The human melanoma cell lines are not reported in the database of commonly misidentified cell lines maintained by ICLAC, were regularly authenticated based on their distinct morphology, and periodically checked for mycoplasma using the Universal Mycoplasma Detection Kit (ATCC). Primary Human Epidermal Melanocytes from adult lightly pigmented donors, (HEMa-LP) were obtained from ThermoFisher Scientific and cultured according to manufacturer instructions in Medium 254 supplemented with Human Melanocyte Growth Supplement-2.

### Transduction of human melanoma cell lines and primary melanocytes

The full-length human SATB2 CDS (IOH46688, Invitrogen) was cloned into pINDUCER20 (Meerbrey et al., 2011) and pLenti CMV Blast DEST (706-1) (Addgene plasmid #17451 was a gift from Eric Campeau & Paul Kaufman) via Gateway recombination using LR clonase II (Invitrogen) according to the manufacturers instructions. Lentiviral particles were produced by cotransfection of 293T cells with sequence verified pINDUCER20-SATB2 or pLentiCMV_SATB2_Blast and packaging plasmids pMD2.G (Addgene plasmid #12259) and psPAX2 (Addgene plasmid #12260, both gifts from Didier Trono), using FuGENE HD (Promega). Viral particles were harvested 48 and 72 hours post-transfection, concentrated by overnight PEG precipitation (Kutner et al., 2009), resuspended in PBS, and stored at −80°C. Human melanoma cell lines and Hema-LP were overlaid with viral particles diluted in DMEM/1x Glutamax with 10% TET System Approved FBS (Clontech) supplemented with 5 μg/ml polybrene (Sigma-Aldrich), for 24 hours at 37°C. 48 hours post-transduction, infected cells were selected with 500 μg/ml G418 (Gibco) for 7 days (pINDUCER20-SATB2), or with 2 μg/ml puromycin (Gibco) (pTK93_Lifeact-mCherry; a gift from Iain Cheeseman, Addgene plasmid #46357) for 3 days, or with 10 μg/ml blasticidin (Gibco) for 3 days (pLentiCMV_SATB2_Blast) replacing selection medium every 48 hours.

### Matrix degradation assay

Oregon green 488-conjungated gelatin covered 12 mm coverslips, or glass bottom 6-well plates were coated as described (Martin et al., 2012). Human melanoma cell lines were induced with doxycycline for 48 hours prior to plating. Human and zebrafish cells were plated at a density of 30,000 cells/well in a 12 well plate, and grown on the fluorescent gelatin for 23-25 hours at 37°C (human cells) or 28°C (zebrafish cells). Cells were fixed in 4% PFA for 10 minutes at room temperature, washed 3 times with PBS, permeabilized with 0.4% Triton-PBS for 4 minutes, followed by 30 minute blocking (10% lab serum, 1% DMSO, 0.1% Tween in PBS). Coverslips were next stained with Alexa 650- or 568-conjungated Phalloidin (1:50) in blocking buffer, overnight at 4°C. Coverslips were washed extensively and nuclei were stained with DAPI, 10 minutes in PBS/0.1% Tween at RT. Coverslips were inversely mounted in Slow Fade Gold (Invitrogen) on a slide, and imaged on a Nikon C2si Laser Scanning Confocal, using ElementsX software. Experiments were seeded in triplicate, in at least 3 independent experiments. Four to six high power (40-60x) areas per coverslip were imaged to determine the fraction of cells with degraded gelatin. An unpaired two-tailed *t*-test was used to compare significance between groups.

### Time-lapse video of fluorescent gelatin degradation after SATB2 induction

Induced human melanoma cell lines were seeded onto an Oregon green 488-conjungated gelatin coated glass bottom 6-well plate, and allowed to adhere for 3 hours at 37°C, prior to imaging. Time-lapse movies were recorded on a Nikon Eclipse Ti Spinning Disk Confocal with a 10X objective (70um, 4um step size), for 16 hrs in 20 minute intervals at 37°C, 5% CO2. Images were processed using Photoshop, ImageJ, or Imaris.

### Proliferation assays

Proliferation rates were determined using the CellTiter-Glo (Promega) luminescent cell viability assay, according to the manufacturer instructions. Cells were seeded at an initial density of 5,000 or 10,000 cells/well (96-well plate), in triplicate or quadruplicate wells, and experiments were repeated at least 3 independent times. An unpaired two-tailed *t*-test was used to compare significance between groups.

### ChlP-seq tumor sample preparation and sequencing

ChIP-seq was performed as previously described (Lee et al., 2006). Zebrafish were euthanized on ice and melanomas were excised, finely minced using a scalpel blade in 5 ml cold PBS in a petri dish, transferred to a 50 ml Falcon tube, and the petri dish was rinsed once with 5 ml PBS. Tumor samples were cross-linked in 1:10 11% formaldehyde solution for 10 minutes at RT, with occasional gentle mixing. Formaldehyde was quenched by addition of 1:20 2.5M glycine. Next, the tumor suspension was mechanically homogenized on ice at the lowest setting for 30 seconds or until no large tumor chunks were visible and passed through a 100 μm filter into a new 50 ml Falcon tube. Cells were spun down at 2500 RPM for 15 minutes, washed twice with PBS, and pellets were flash frozen and stored at −80°C, or subjected to chromatin immunoprecipitation with anti-SATB2 (sc-81376, Santa Cruz), anti-Histone H3 acetyl K27 (ab4729, Abcam) or anti-Histone H3 tri methyl K9 (ab8898, Abcam) antibodies. 10 μl of input DNA and the entire volume of ChIP DNA samples were prepared for sequencing. Libraries were prepared using the NEBNext Multiplex Oligos Kit (18-cycle PCR enrichment step), mixed in equal quantities (2-10 nM), and run on an Illumina Hi-Seq2000.

### ChlP-sequencing bioinformatics

All ChIP-Seq datasets were aligned to build version danRer7 of the zebrafish genome using Bowtie2 (version 2.2.1) (Langmead and Salzberg, 2012) with the following parameters: --end-to-end, -N0, -L20. We used the MACS2 version 2.1.0 (Zhang et al., 2008) peak finding algorithm to identify regions of ChIP-Seq peaks, with a *q*-value threshold of enrichment of 0.05 for all datasets (Langmead and Salzberg, 2012). Genome track images were generated using the UCSC browser. SATB2-bound loci were determined by overlapping all regions 3 kb upstream of the transcriptional start site (TSS), gene body (GB) and 3 kb downstream of the transcriptional end site (TES) of all transcripts in the danRer7 assembly, with all significantly bound SATB2-peaks (*P*<10^-7^). SATB2 bound peaks were compared to SATB2 enhancers peaks to determine the SATB2-bound enhancers. GREAT (Hiller et al., 2013) version 2.0.2 was used for GO-term enrichment analysis of SATB2-associated loci and SATB2-bound enhancers, using the ‘Basal plus extension’ genomic region association (proximal 10 kb upstream of TSS, 5 kb downstream of TSS, distal up to 100 kb) to associate genomic regions with nearby genes. The list of SATB2-bound genes was converted to human orthologs using DIOPT – DRSC Integrative Ortholog Prediction Tool (Hu et al., 2011) returning only the best matching ortholog. Data sets are deposited to the GEO Gene Expression Omnibus, accession number GSE77923 (http://www.ncbi.nlm.nih.gov/geo/query/acc.cgi?token=qdibymmypdktrsz&acc=GSE77923).

### RNA-sequencing bioinformatics

RNA-seq was performed on three biological replicates of primary zebrafish melanoma tumors that were excised from MCR:EGFP (control) and MCR:SATB2 overexpressing *Tg(BRAF^ve00E^);p53^-/-^; mitf^-/-^* zebrafish. Zebrafish were euthanized on ice, melanomas were isolated and mechanically homogenized in RTL buffer (Qiagen) containing β-mercaptoethanol. Tumor lysates were transferred onto a QiaShredder column (Qiagen), and RNA isolation was performed using the RNA Micro Plus kit (Qiagen) according to the manufacturers instruction. RNA quality was determined using a Nanodrop and Fragment Analyzer (Advanced Analytical). Total RNA was depleted of ribosomal RNA with the Ribo-Zero Gold kit (Epicentre). Ribosome-depleted RNA was used to create multiplexed RNA-seq libraries (NEBNext Ultra, NEB) according to manufacturer’s instructions. Samples with an average DNA input size of 300 base pairs were sequenced on an Illumina Hi-Seq2000 sequencer with 2 × 100-bp paired-end reads. Quality control of RNA-Seq datasets was performed by FastQC29 and Cutadapt30 to remove adaptor sequences and low quality regions. The high-quality reads were aligned to UCSC build danRer7 of zebrafish genome using Tophat31 2.0.11 without novel splicing form calls. Transcript abundance and differential expression were calculated with Cufflinks32 2.2.1. FPKM values were used to normalize and quantify each transcript; the resulting list of differential expressed genes are filtered by log(2) fold change > 1.5 and a *P*-value < 0.05. Human orthologs were predicted using DIOPT33. Data sets are deposited to the GEO Gene Expression Omnibus, accession number GSE77923. RNA-seq was performed on three biological replicates of SKMEL2 transduced with pINDUCER20-SATB2 after 48 hours of induction with 2ug/mL of doxycycline as outlined above, except for the mechanical homogenization. Pathway analyses were conducted using GSEA (v4.0) (Subramanian et al., 2005) and IPA (Qiagen). Pathway and gene list overlaps were plotted using BioVenn (Hulsen et al., 2008).

### Statistics

Comparison of Kaplan-Meier survival curves was performed by a log-rank (Mantel-Cox) test. Statistical difference in qPCR expression analysis between large groups of MCR:EGFP and MCR:SATB2 primary tumors, and difference in estimated fraction of tumor propagating cells between MCR:EGFP and MCR:SATB2 donors were determined by an unpaired two-tailed Mann-Whitney test. The rest of the statistics were performed with an unpaired two-tailed t-test. An F-test was used to determine similar variation between compared groups. Graphs show the median with s.e.m. No statistical methods were used to predetermine sample size. Experiments to quantify proliferation by PH3 immunohistochemistry, and cells with degraded 488-conjungated gelatin were scored blindly. Irradiated *casper* zebrafish were randomized between transplantation groups. All statistical analyses were performed with GraphPad Prism. NS, not significant, *P* > 0.05; *\*P* ≤ 0.05; *\*\*P* ≤ 0.01; *** *P* ≤ 0.001; **** *P* ≤ 0.0001

## Supplemental Figure

**Supplementary Figure 1:**
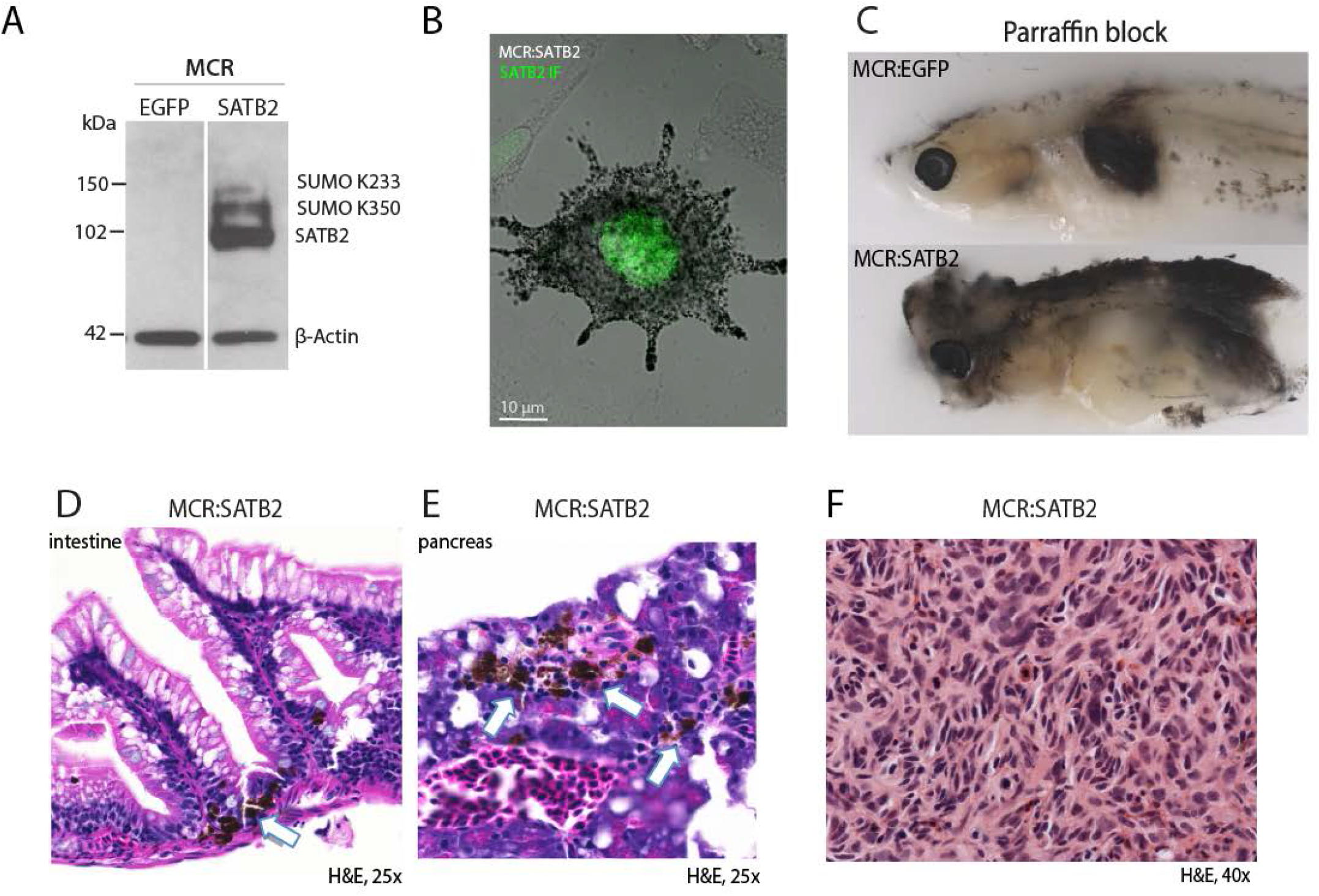
SATB2 overexpression leads to invasive melanoma with organ involvement. (A) Western Blot and (B) immunofluorescence validation of human SATB2 protein expression in MCR:SATB2 zebrafish primary tumors. (C) Cross section through zebrafish mounted in a paraffin block. Compared to the MCR:EGFP-injected zebrafish which has a pigmented tumor with confined borders, in the MCR:SATB2-injected zebrafish dispersed pigmented tumor cells can be observed throughout the body. MCR:SATB2-injected zebrafish have increase organ involvement, as shown by pigmented melanoma cells (white arrow) in the (D) intestine, and (E) pancreas. (F) Histological aspect of MCR:SATB2 tumors illustrating spindle morphology of melanoma cells.

**Supplementary Figure 2:**
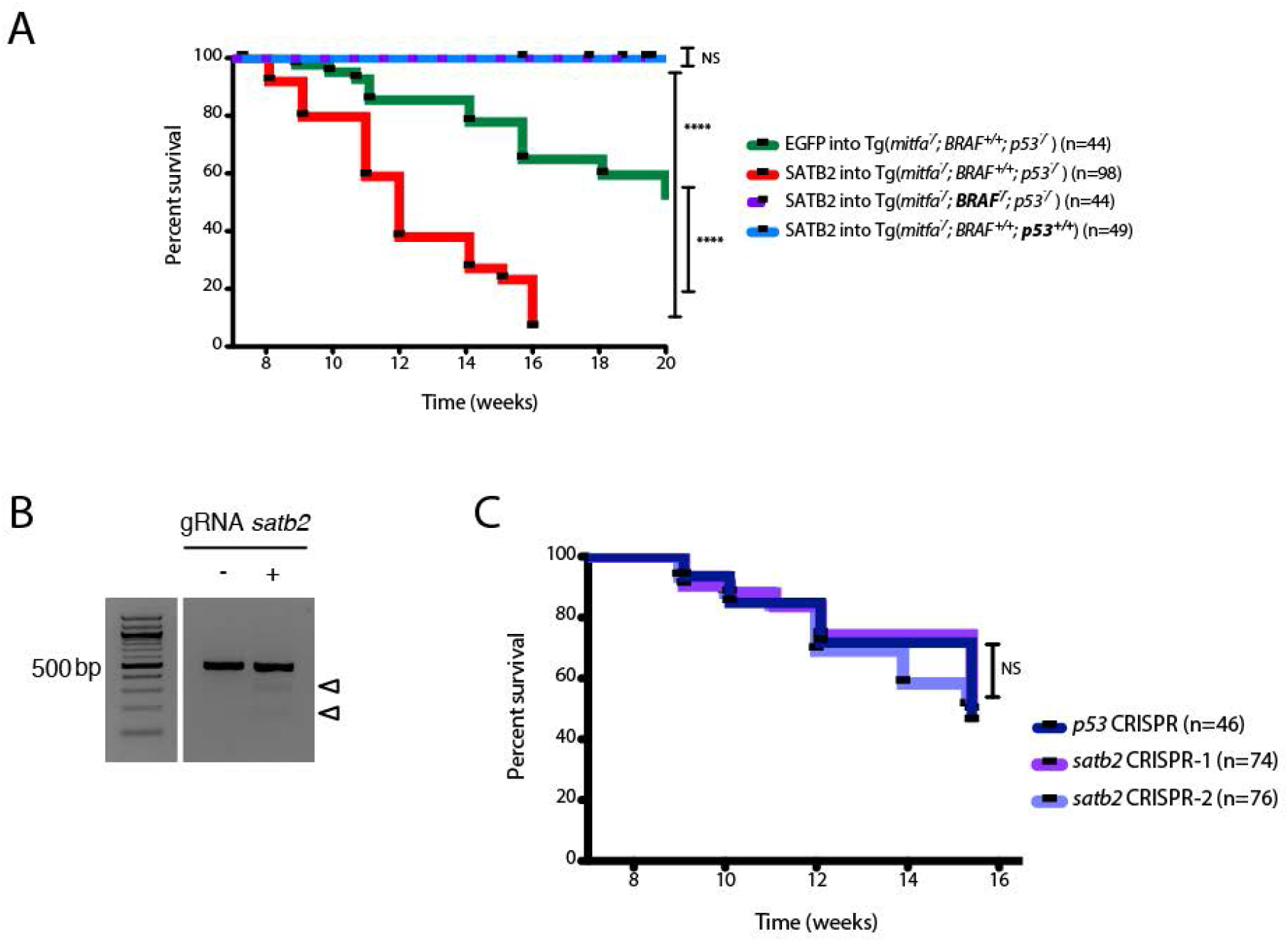
SATB2 is neither sufficient nor necessary for melanoma initiation in zebrafish. (A) Kaplan-Meier survival curves showing that MCR:SATB2 acceleration is dependent on *BRAF^V600E^* and *p53^-/-^.* Plotted: MCR:EGFP (n=44) and MCR:SATB2 (n=98) into *Tg(mitfa:BRAF^V600E^)*; p53^-/-^; mitfa^-/-^ compared to MCR:SATB2 into *BRAF^-/-^; p53^-/-^* (n= 44) and MCR:SATB2 into *BRAF^+/+^;p53^+/+^* (n= 49) (logrank Mantel-Cox). (B) T7E1 mutagenesis assay at the CRISPR target site in the *satb2* gene. The assay was performed on genomic DNA from 2 dpf embryos injected at the one-cell stage with *Cas9* mRNA and either a control gRNA (left) or a gRNA against *satb2* (right). Cleavage bands (arrowheads, expected sizes 167 and 303 bp) indicate efficient mutagenesis at the target site. (C) Kaplan-Meier survival curves for *p53* CRISPR (n-46), *satb2* CRISPR-1 (n=74), *satb2* CRISPR-2 (n=76) injected into [Tg(*mitfa:BRAF^V600E^*); *p53^-/-^*; *mitfa^-/-^*] reveal *satb2* is dispensable for melanoma initiation. Logrank NS; not significant, \*\*\*\**P*<0.0001.

**Supplementary Figure 3:**
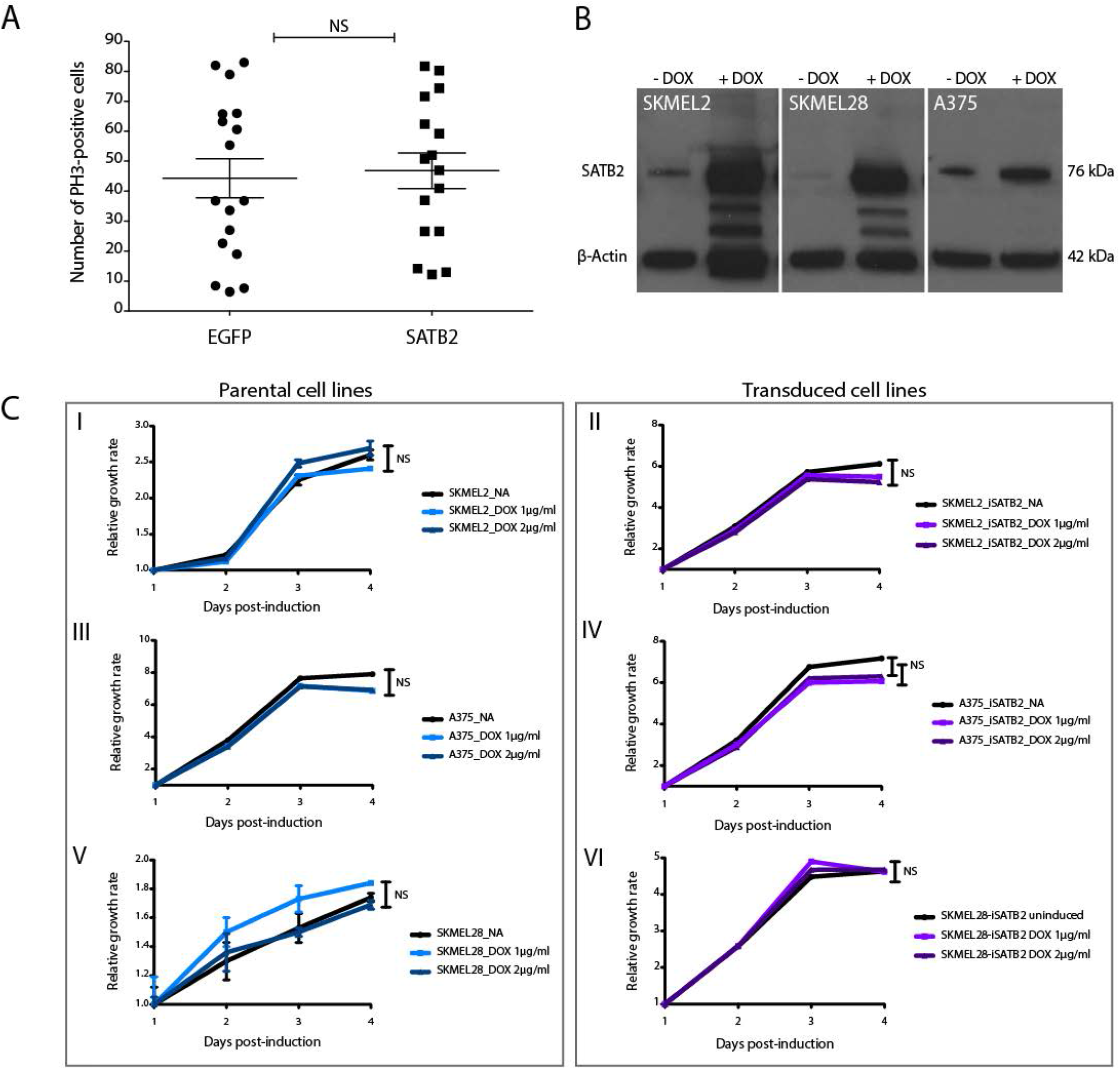
SATB2 overexpression does not affect proliferation. (A) Proliferation rates in primary MCR:EGFP (n=17) and MCR:SATB2 (n=16) tumors are not statistically different (unpaired Student’s *t*-test), as determined by PH3 immunohistochemistry. (B) Western blot for SATB2 after DOX-induction in SKMEL2, SKMEL28 and A375 human melanoma cell lines. β-Actin was used as a loading control. The two lower bands in SKMEL2 and SKMEL28 are likely degradation products of SATB2 due to the overexpression. (C) Induction of SATB2 does not affect proliferation rates between parental and transduced cell lines, as determined by CellTiter-GLO (NS; not significant).

**Supplementary Figure 4:**
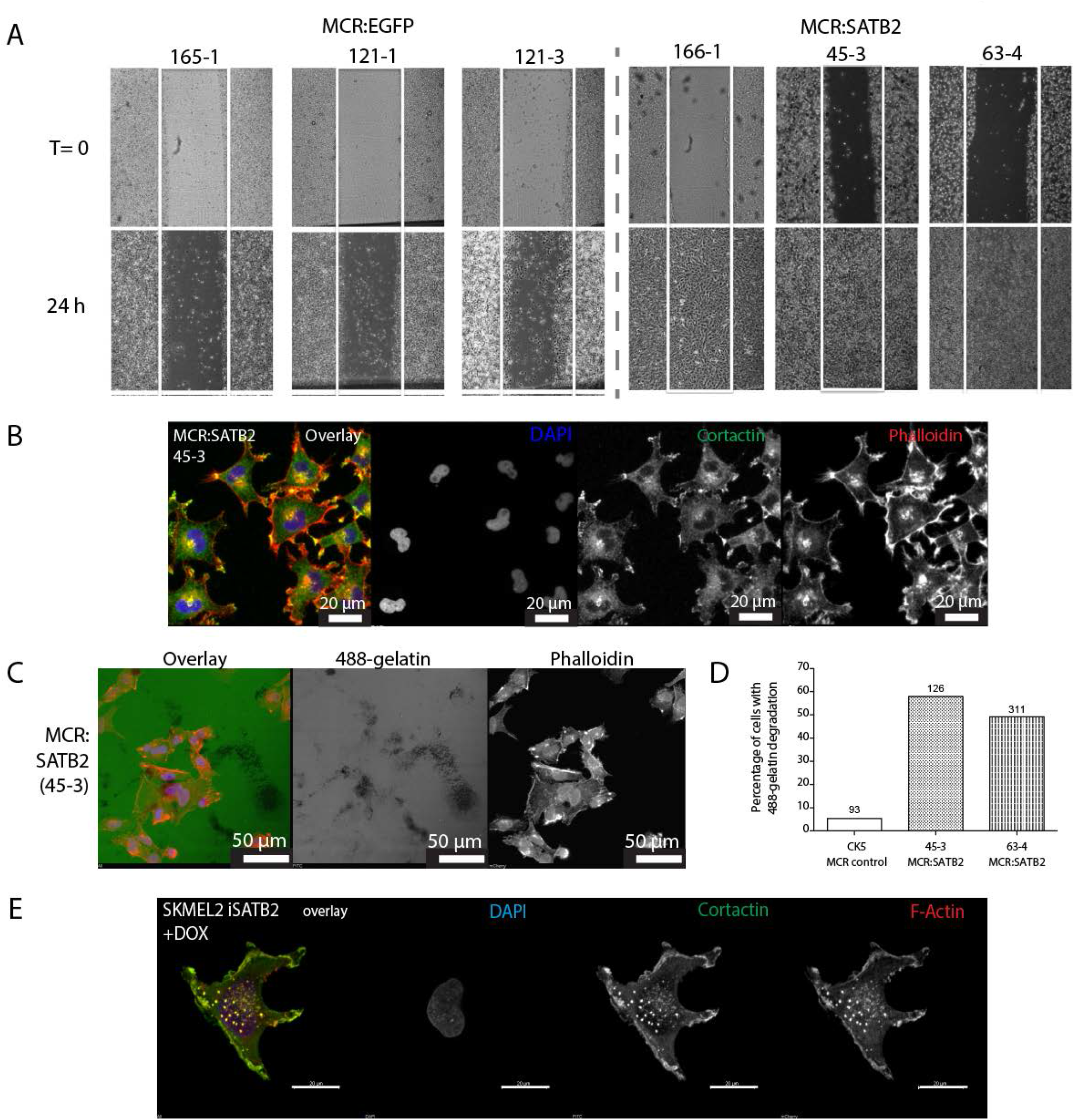
SATB2 overexpression induces EMT, increased migration and invadopodia formation in zebrafish melanoma. (A) Scratch assays on cultured MCR tumors show increased migration in MCR:SATB2 tumors. (B) Confocal analysis of immunohistochemistry of cultured MCR:SATB2 tumor shows co-localization of Cortactin (green) and F-actin (phalloidin, red), suggesting the formation of invadopodia. DAPI nuclear stain is shown in blue. (C) MCR:SATB2 (45-3) cells show increased Oregon green 488-conjungated gelatin degradation 24-25 hours post-seeding. Scale bar is 50 μm. (D) Percentage of zebrafish melanoma cells (n) with degraded 488-conjungated gelatin 24 hours post-seeding, observed in MCR (CK5, n=93), MCR:SATB2 (45-3, n=126) and MCR:SATB2 (63-4, n=311) cell lines. (E) Immunofluorescence showing colocalization of F-actin and Cortactin in SKMEL2 overexpressing SATB2 in the presence of doxycycline (48h induction).

**Supplementary Figure 5:**
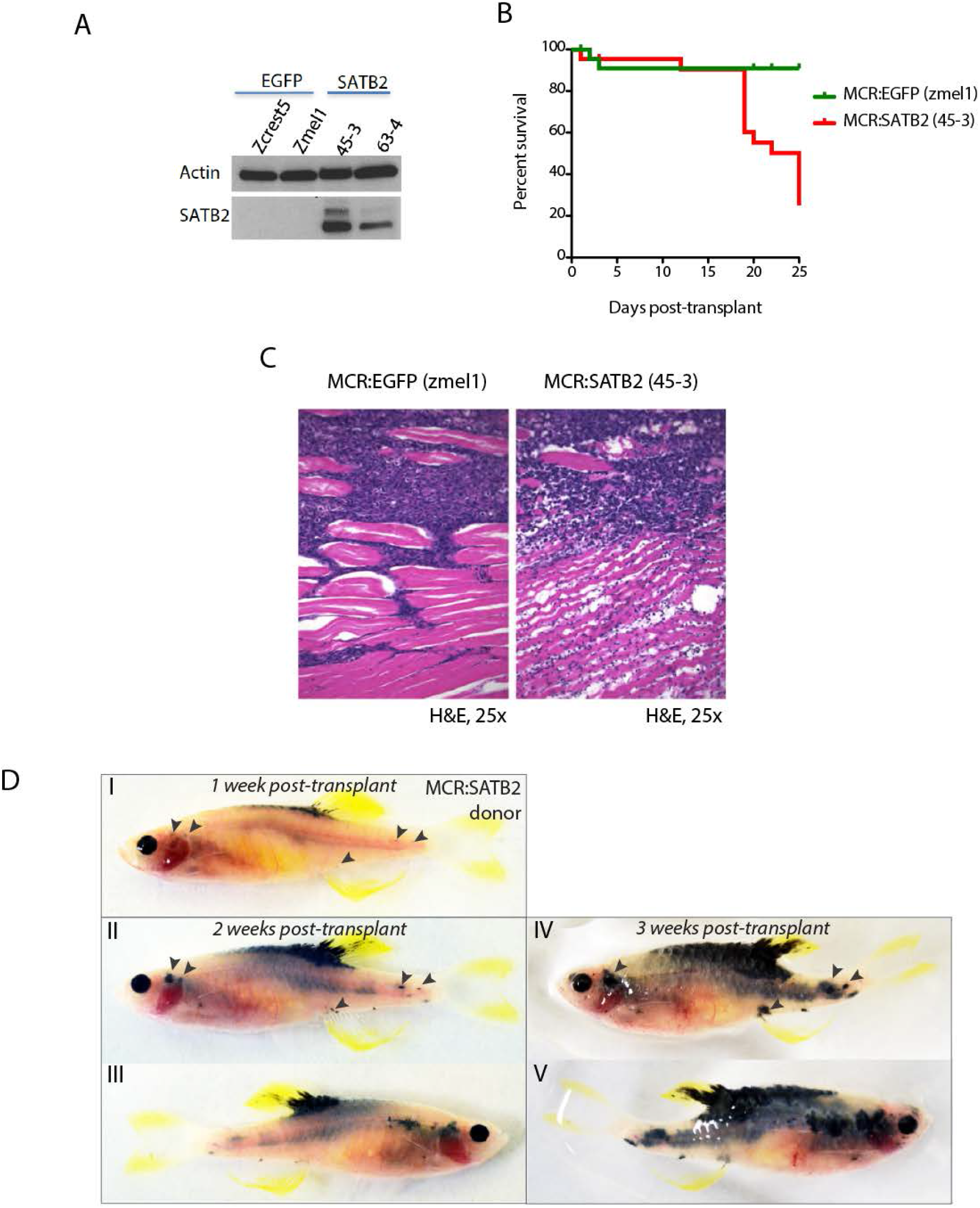
MCR:SATB2 primary tumors and long term cultures have increase invasion potential in vivo. (A) Western blot of SATB2 showing retained expression in long term *in vitro* cultures derived from MCR:SATB2 tumors. Actin loading control is shown. (B) Overall survival and (C) histology of irradiated *casper* recipients transplanted with MCR:EGFP Zmel1 (n=31) or MCR:SATB2 45-3 (n=31) zebrafish long term *in vitro* cultures. (D) Weekly prospective imaging of primary MCR:SATB2 tumor allograft in irradiated *casper* recipient. Arrows indicate distant metastasis from injection site. Related to Figure 2H.

**Supplementary Figure 6:**
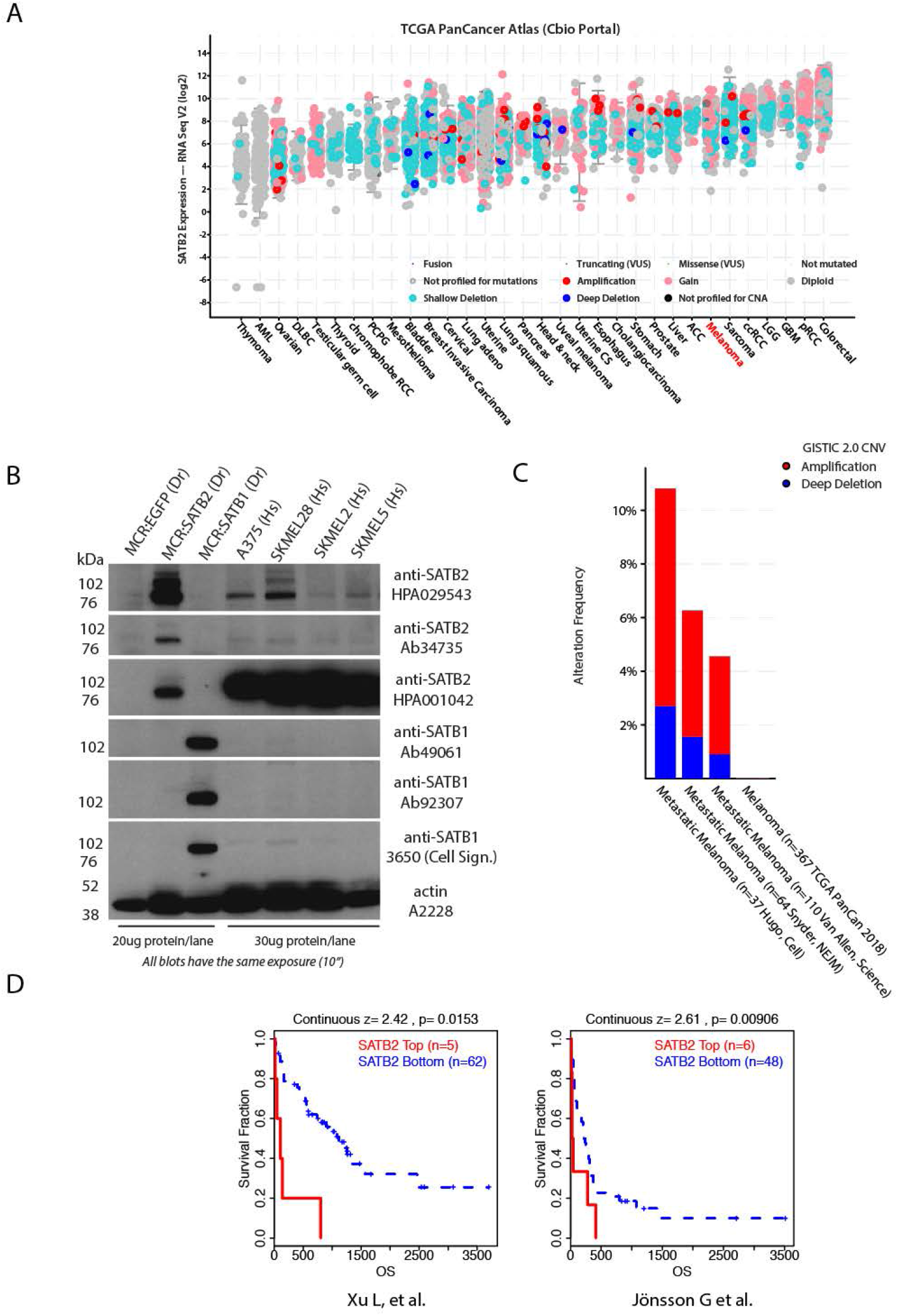
Analysis of SATB2 expression, amplification and correlation with survival in human melanoma. (A) Analysis of SATB2 expression across tumor types in the TCGA PanCancer dataset using cBio portal (Gao et al., 2013). Melanoma (highlighted in red) SATB2 mRNA levels are comparable to other solid tumors where SATB2 has been established to play a role (i.e. RCC, Sarcoma, CRC). (B) Western blot of SATB2 and SATB1 with a panel of antibodies showing protein expression in a panel of human melanoma cell lines and zebrafish MCR:EGFP, MCR:SATB2 and MCR:SATB1 control tumors. Actin loading control is shown. (C) Frequency of SATB2 amplification in red across multiple available melanoma genomic studies (Consortium, 2020; Hugo et al., 2015; Snyder et al., 2014; Van Allen et al., 2014). (D) Correlation between SATB2 expression and melanoma metastasis-related overall survival in two independent publicly available dataset (GSE8401 and GSE22153) of melanoma followed for metastatic progression risk. Continuous expression Z-score optimized plots from TIDE Harvard Portal http://tide.dfci.harvard.edu/ (Fu et al., 2020).

**Supplementary Figure 7:**
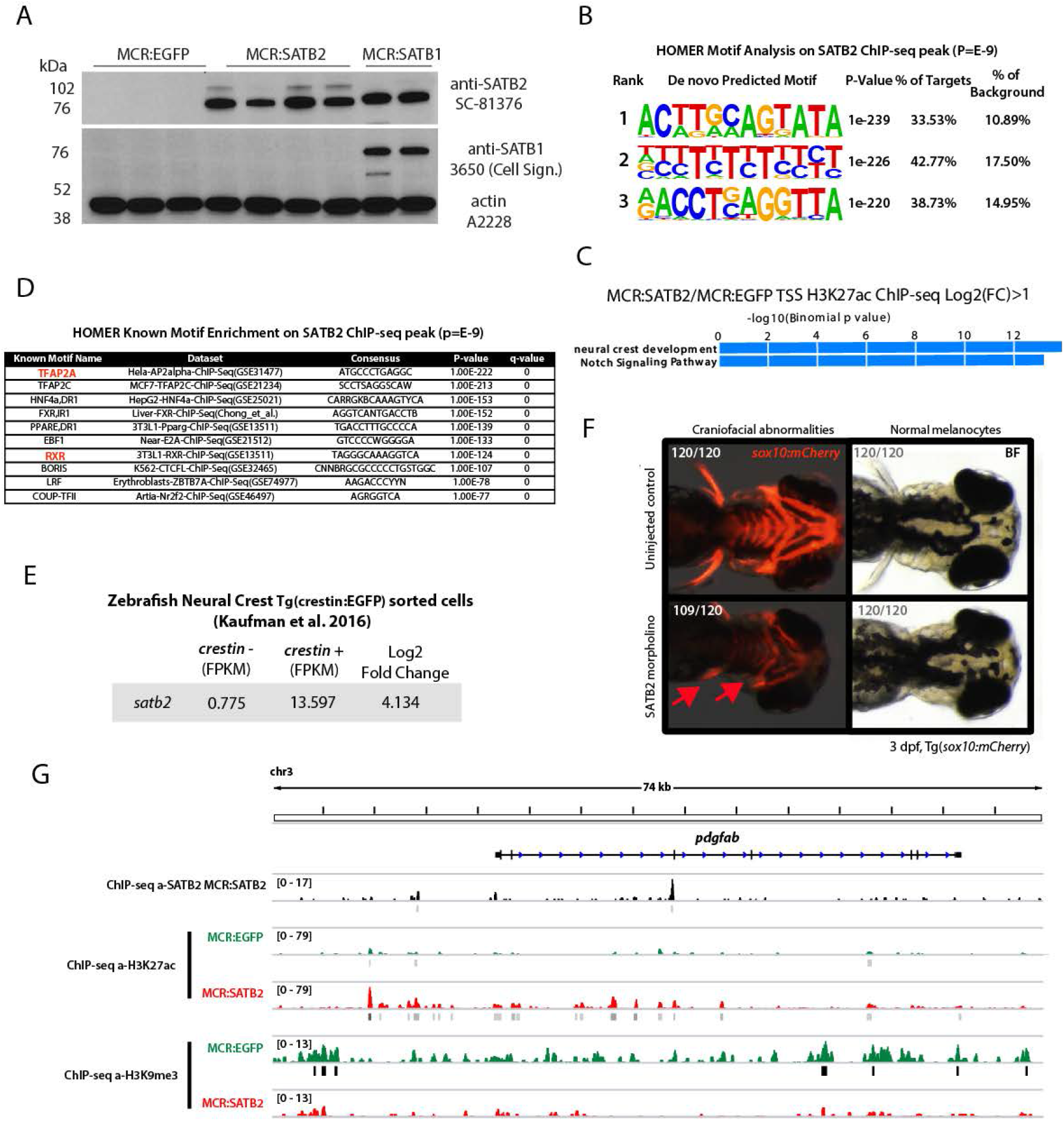
Analysis of SATB2 expression and chromatin effect in human and zebrafish melanoma. (A) Western blot analysis in MCR:EGFP, MCR:SATB2, and MCR:SATB1 zebrafish tumors of SATB2 expression and antibody cross reactivity with human SATB1 (in MCR:SATB1) and lack of detection of endogenous zebrafish satb2 and satb1 (in MCR:EGFP). The SC-81376 antibody was used for anti-SATB2 ChIP-seq in MCR:SATB2 tumors, since the other anti-SATB2 antibodies shown in Supplementary Figure 6B did not work in immunoprecipitation. (B) HOMER motif prediction analysis of significant peaks (*p*<E-9) from anti-SATB2 ChIP-seq enrichment over input in MCR:SATB2 tumors showing the top motif to be AT-rich, as expected. (C) Go-term enrichment analysis of genome wide H3K27Ac peaks enriched in MCR:SATB2 over MCR:EGFP Around TSS Log2FC>1(GREAT analysis). (D) HOMER known motif enrichment analysis on anti-SATB2 ChIP-seq peaks (*p*<E-9) in MCR:SATB2. (E) RNA-seq analysis of *satb2* expression in sorted *crestin:EGFP^+^* developing neural crest cells (data source (Kaufman et al., 2016)). (F) Knock-down of *satb2* in the transgenic neural crest reporter line *Tg(sox10:mCherry)* using a validated morpholino for *satb2.* While *satb2* knock-down did not affect melanocyte development (0/120), *satb2* morphants displayed severe cranio-facial abnormalities as observed by overall reduced *sox10* reporter expression, reduced pectoral fins (red arrows), and head size (109/120). (G) Example analysis of the chromatin state of a direct SATB2 target showing the *pdgfab* locus. ChIP-Seq track in zebrafish MCR:SATB2 for SATB2 (in black). ChIP-seq tracks for H3K27Ac and H3K9me3 in primary tumors MCR:SATB2 (red) and MCR:EGFP controls (green).

**Supplementary Figure 8:**
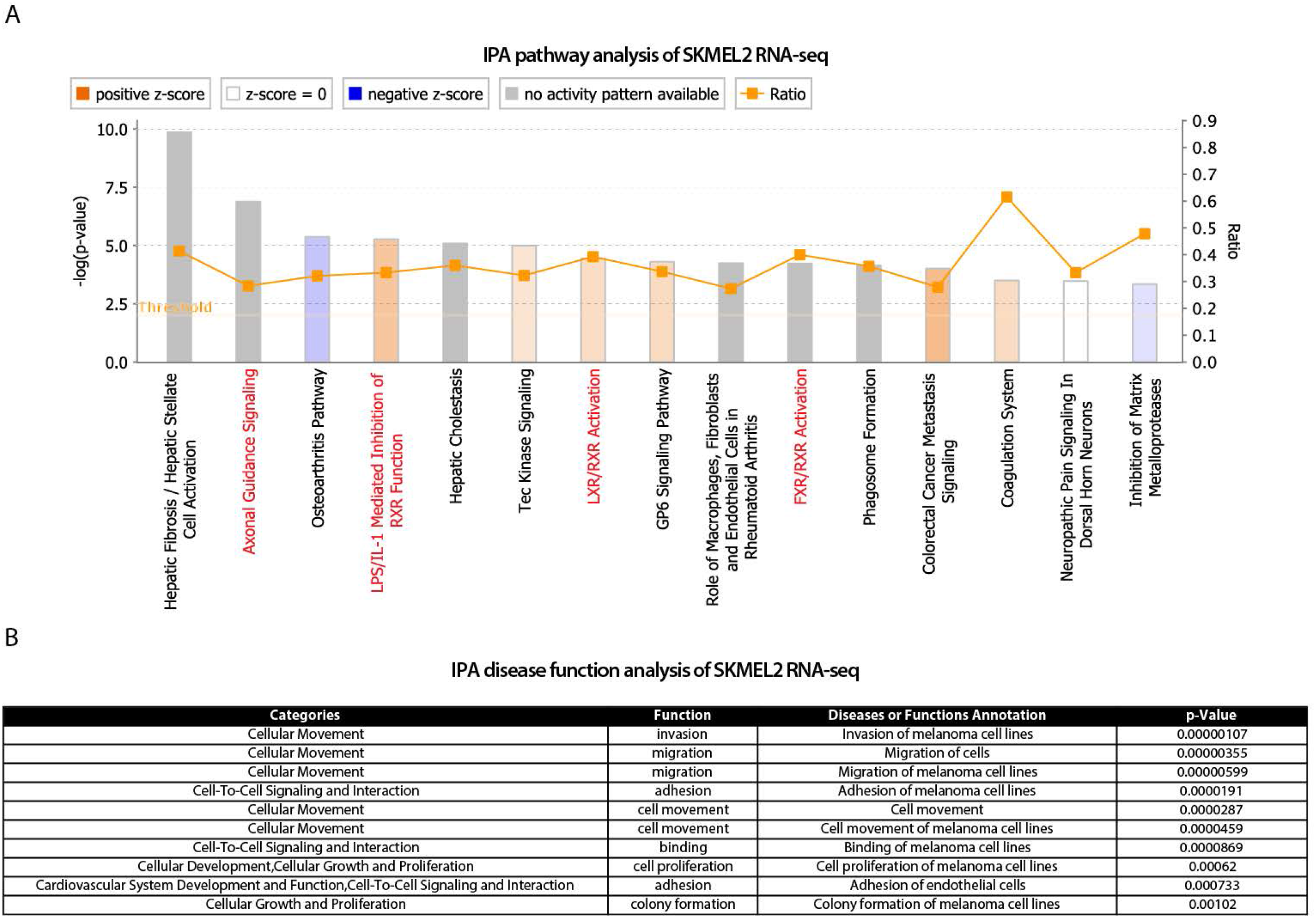
IPA Analysis of significant differentially expressed genes in SKMEL2 iSATB2. IPA analysis on (A) Pathways and (B) disease function on SKMEL2 iSATB2. Related to Figure 4B-C.

**Supplementary Table 1**: Schematic of screened pools of Epigenetic factors and source plasmid references

**Supplementary Table 2**: (Sheet 1)Differential gene expression list of primary MCR:SATB2 vs MCR:EGFP tumors by RNA-seq. (Sheet 2) Genes confirmed by qPCR on a large panel of primary MCR:SATB2 tumors versus MCR:EGFP tumors. Red is upregulated, green is downregulated. (Sheet 3) Best human ortholog prediction with DIOPT. (Sheet 4) MCR tumors RNA-seq data matched with human orthologs. (Sheet 5) Input for GSEA analysis on MCR tumors. (Sheet 6) Input for IPA analysis on MCR tumors. (Sheet 7) SATB2 bound genes based on SATB2 ChIP-seq in MCR:SATB2 tumor. (Sheet 8) DIOPT human ortholog prediction of SATB2 bound targets, and (Sheet 9) humanized SATB2 bound target list used for overlap in Biovenn with published TFAP2A active targets in Figure 3D. (Sheet 10) Overlap of significant genes with with ChIP-seq to determine SATB2-bound and SATB2-associated loci that are misexpressed.

**Supplementary Table 3: SATB2-associated genes that are neural crest related** (Sheet 1) SATB2-associated genes, determined by using GREAT genomic region association ‘Basal plus extension’ (proximal 10 kb upstream of TSS, 5 kb downstream of TSS, distal up to 100 kb). (Sheet 2) Neural crest associated genes. (Sheet 3) SATB2/neural crest-associated genes. (Sheet 4) Human predicted orthologs (DIOPT) of SATB2-Bound genes (TSS+ gene body +/− 3kb).

**Supplementary Table 4: RNA-seq of SKMEL2 TetOn SATB2 48hours +/− doxycycline**

(Sheet 1) Genes with a log(2) fold change > 1.0 and a *P*-value < 0.05. (Sheet 2) Unfiltered RNA-seq expression data.

**Supplementary Table 5: qRT-PCR primers**

**Supplemental Video 1**: Zebrafish MCR:EGFP (unpigmented) and MCR:SATB2 (pigmented) melanoma phenotype

**Supplemental Video 2**: Time-lapse video recording of Oregon Green 488-labeled gelatin degradation upon SATB2 induction in human melanoma cell line A375

**Supplemental Video 3**: MCR control primary tumor transplant into transparent *casper* zebrafish, 3.5 weeks post-transplant

**Supplemental Video 4**: MCR:SATB2 primary tumor transplant into transparent *casper* zebrafish, 3.5 weeks post-transplant

## References

Ablain, J., Xu, M., Rothschild, H., Jordan, R. C., Mito, J. K., Daniels, B. H., … Yeh, I. (2018). Human tumor genomics and zebrafish modeling identify SPRED1 loss as a driver of mucosal melanoma. Science, 362(6418), 1055–1060. doi:10.1126/science.aau6509

Ahmed, F., & Haass, N. K. (2018). Microenvironment-Driven Dynamic Heterogeneity and Phenotypic Plasticity as a Mechanism of Melanoma Therapy Resistance. Front Oncol, 8, 173. doi:10.3389/fonc.2018.00173

Ahn, H. J., Park, Y., Kim, S., Park, H. C., Seo, S. K., Yeo, S. Y., & Geum, D. (2010). The expression profile and function of Satb2 in zebrafish embryonic development. Mol Cells, 30(4), 377–382. doi:10.1007/s10059-010-0128-6

Bailey, C. M., Morrison, J. A., & Kulesa, P. M. (2012). Melanoma revives an embryonic migration program to promote plasticity and invasion. Pigment Cell Melanoma Res, 25(5), 573–583. doi:10.1111/j.1755-148X.2012.01025.x

Bajpai, R., Chen, D. A., Rada-Iglesias, A., Zhang, J., Xiong, Y., Helms, J., … Wysocka, J. (2010). CHD7 cooperates with PBAF to control multipotent neural crest formation. Nature, 463(7283), 958–962. doi:10.1038/nature08733

Betancur, P., Bronner-Fraser, M., & Sauka-Spengler, T. (2010). Assembling neural crest regulatory circuits into a gene regulatory network. Annu Rev Cell Dev Biol, 26, 581–603. doi:10.1146/annurev.cellbio.042308.113245

Birkeland, E., Zhang, S., Poduval, D., Geisler, J., Nakken, S., Vodak, D., … Lonning, P. E. (2018). Patterns of genomic evolution in advanced melanoma. Nat Commun, 9(1), 2665. doi:10.1038/s41467-018-05063-1

Boshuizen, J., Vredevoogd, D. W., Krijgsman, O., Ligtenberg, M. A., Blankenstein, S., de Bruijn, B., … Peeper, D. S. (2020). Reversal of pre-existing NGFR-driven tumor and immune therapy resistance. Nat Commun, 11(1), 3946. doi:10.1038/s41467-020-17739-8

Boumahdi, S., & de Sauvage, F. J. (2020). The great escape: tumour cell plasticity in resistance to targeted therapy. Nat Rev Drug Discov, 19(1), 39–56. doi:10.1038/s41573-019-0044-1

Cancer Genome Atlas, N. (2015). Genomic Classification of Cutaneous Melanoma. Cell, 161(7), 1681–1696. doi:10.1016/j.cell.2015.05.044

Ceol, C. J., Houvras, Y., Jane-Valbuena, J., Bilodeau, S., Orlando, D. A., Battisti, V., … Zon, L. I. (2011). The histone methyltransferase SETDB1 is recurrently amplified in melanoma and accelerates its onset. Nature, 471(7339), 513–517. doi:10.1038/nature09806

Consortium, I. T. P.-C. A. o. W. G. (2020). Pan-cancer analysis of whole genomes. Nature, 578(7793), 82–93. doi:10.1038/s41586-020-1969-6

Dang, M., Henderson, R. E., Garraway, L. A., & Zon, L. I. (2016). Long-term drug administration in the adult zebrafish using oral gavage for cancer preclinical studies. Disease models & mechanisms, 9(7), 811–820. doi:10.1242/dmm.024166

Ding, L., Kim, M., Kanchi, K. L., Dees, N. D., Lu, C., Griffith, M., … Weber, J. S. (2014). Clonal architectures and driver mutations in metastatic melanomas. PLoS One, 9(11), e111153. doi:10.1371/journal.pone.0111153

Ekpe-Adewuyi, E., Lopez-Campistrous, A., Tang, X., Brindley, D. N., & McMullen, T. P. (2016). Platelet derived growth factor receptor alpha mediates nodal metastases in papillary thyroid cancer by driving the epithelial-mesenchymal transition. Oncotarget, 7(50), 83684–83700. doi:10.18632/oncotarget.13299

Fallahi-Sichani, M., Becker, V., Izar, B., Baker, G. J., Lin, J. R., Boswell, S. A., … Sorger, P. K. (2017). Adaptive resistance of melanoma cells to RAF inhibition via reversible induction of a slowly dividing de-differentiated state. Mol Syst Biol, 13(1), 905. doi:10.15252/msb.20166796

Fazio, M., van Rooijen, E., Mito, J. K., Modhurima, R., Weiskopf, E., Yang, S., & Zon, L. I. (2020). Recurrent co-alteration of HDGF and SETDB1 on chromosome 1q drives cutaneous melanoma progression and poor prognosis. Pigment Cell Melanoma Res. doi:10.1111/pcmr.12937

Gan, X., Jiang, J., Wu, G., Chen, D., Liao, D., & Li, M. (2017). SATB2 induces stem-like properties and promotes epithelial-mesenchymal transition in hepatocellular carcinoma. Int J Clin Exp Pathol, 10(12), 11932–11940. Retrieved from https://www.ncbi.nlm.nih.gov/pubmed/31966558

Gupta, P. B., Kuperwasser, C., Brunet, J. P., Ramaswamy, S., Kuo, W. L., Gray, J. W., … Weinberg, R. A. (2005). The melanocyte differentiation program predisposes to metastasis after neoplastic transformation. Nat Genet, 37(10), 1047–1054. doi:10.1038/ng1634

Gyorgy, A. B., Szemes, M., de Juan Romero, C., Tarabykin, V., & Agoston, D. V. (2008). SATB2 interacts with chromatin-remodeling molecules in differentiating cortical neurons. Eur J Neurosci, 27(4), 865–873. doi:10.1111/j.1460-9568.2008.06061.x

Hassan, M. Q., Gordon, J. A., Beloti, M. M., Croce, C. M., van Wijnen, A. J., Stein, J. L., … Lian, J. B. (2010). A network connecting Runx2, SATB2, and the miR-23a~27a~24-2 cluster regulates the osteoblast differentiation program. Proc Natl Acad Sci U S A, 107(46), 19879–19884. doi:10.1073/pnas.1007698107

Heilmann, S., Ratnakumar, K., Langdon, E., Kansler, E., Kim, I., Campbell, N. R., … White, R. M. (2015). A quantitative system for studying metastasis using transparent zebrafish. Cancer Res. doi:10.1158/0008-5472.CAN-14-3319

Hugo, W., Shi, H., Sun, L., Piva, M., Song, C., Kong, X., … Lo, R. S. (2015). Non-genomic and Immune Evolution of Melanoma Acquiring MAPKi Resistance. Cell, 162(6), 1271–1285. doi:10.1016/j.cell.2015.07.061

Hugo, W., Zaretsky, J. M., Sun, L., Song, C., Moreno, B. H., Hu-Lieskovan, S., … Lo, R. S. (2016). Genomic and Transcriptomic Features of Response to Anti-PD-1 Therapy in Metastatic Melanoma. Cell, 165(1), 35–44. doi:10.1016/j.cell.2016.02.065

Iwanaga, R., Truong, B. T., Hsu, J. Y., Lambert, K. A., Vyas, R., Orlicky, D., … Artinger, K. B. (2020). Loss of prdm1a accelerates melanoma onset and progression. Mol Carcinog, 59(9), 1052–1063. doi:10.1002/mc.23236

Johannessen, C. M., Johnson, L. A., Piccioni, F., Townes, A., Frederick, D. T., Donahue, M. K., … Garraway, L. A. (2013). A melanocyte lineage program confers resistance to MAP kinase pathway inhibition. Nature, 504(7478), 138–142. doi:10.1038/nature12688

Kapp, F. G., Perlin, J. R., Hagedorn, E. J., Gansner, J. M., Schwarz, D. E., O’Connell, L. A., … Zon, L. I. (2018). Protection from UV light is an evolutionarily conserved feature of the haematopoietic niche. Nature, 558(7710), 445–448. doi:10.1038/s41586-018-0213-0

Kaufman, C. K., Mosimann, C., Fan, Z. P., Yang, S., Thomas, A. J., Ablain, J., … Zon, L. I. (2016). A zebrafish melanoma model reveals emergence of neural crest identity during melanoma initiation. Science, 351(6272), aad2197. doi:10.1126/science.aad2197

Kikuiri, T., Mishima, H., Imura, H., Suzuki, S., Matsuzawa, Y., Nakamura, T., … Yoshiura, K. I. (2018). Patients with SATB2-associated syndrome exhibiting multiple odontomas. Am J Med Genet A, 176(12), 2614–2622. doi:10.1002/ajmg.a.40670

Li, F. Z., Dhillon, A. S., Anderson, R. L., McArthur, G., & Ferrao, P. T. (2015). Phenotype switching in melanoma: implications for progression and therapy. Front Oncol, 5, 31. doi:10.3389/fonc.2015.00031

Li, P., White, R. M., & Zon, L. I. (2011). Transplantation in zebrafish. Methods Cell Biol, 105, 403–417. doi:10.1016/B978-0-12-381320-6.00017-5

Maheswaran, S., & Haber, D. A. (2015). Cell fate: Transition loses its invasive edge. Nature, 527(7579), 452–453. doi:10.1038/nature16313

Mansour, M. A., Asano, E., Hyodo, T., Akter, K. A., Takahashi, M., Hamaguchi, M., & Senga, T. (2015). Special AT-rich sequence-binding protein 2 suppresses invadopodia formation in HCT116 cells via palladin inhibition. Exp Cell Res, 332(1), 78–88. doi:10.1016/j.yexcr.2014.12.003

Marine, J. C., Dawson, S. J., & Dawson, M. A. (2020). Non-genetic mechanisms of therapeutic resistance in cancer. Nat Rev Cancer. doi:10.1038/s41568-020-00302-4

Martin, K. H., Hayes, K. E., Walk, E. L., Ammer, A. G., Markwell, S. M., & Weed, S. A. (2012). Quantitative measurement of invadopodia-mediated extracellular matrix proteolysis in single and multicellular contexts. J Vis Exp(66), e4119. doi:10.3791/4119

Mayor, R., & Theveneau, E. (2013). The neural crest. Development, 140(11), 2247–2251. doi:10.1242/dev.091751

McConnell, A. M., Mito, J. K., Ablain, J., Dang, M., Formichella, L., Fisher, D. E., & Zon, L. I. (2018). Neural crest state activation in NRAS driven melanoma, but not in NRAS-driven melanocyte expansion. Dev Biol. doi:10.1016/j.ydbio.2018.05.026

McKenna, W. L., Ortiz-Londono, C. F., Mathew, T. K., Hoang, K., Katzman, S., & Chen, B. (2015). Mutual regulation between Satb2 and Fezf2 promotes subcerebral projection neuron identity in the developing cerebral cortex. Proc Natl Acad Sci U S A, 112(37), 11702–11707. doi:10.1073/pnas.1504144112

Murphy, D. A., & Courtneidge, S. A. (2011). The ‘ins’ and ‘outs’ of podosomes and invadopodia: characteristics, formation and function. Nat Rev Mol Cell Biol, 12(7), 413–426. doi:10.1038/nrm3141

Murphy, D. A., Diaz, B., Bromann, P. A., Tsai, J. H., Kawakami, Y., Maurer, J., … Courtneidge, S. A. (2011). A Src-Tks5 pathway is required for neural crest cell migration during embryonic development. PLoS One, 6(7), e22499. doi:10.1371/journal.pone.0022499

Naik, R., & Galande, S. (2019). SATB family chromatin organizers as master regulators of tumor progression. Oncogene, 38(12), 1989–2004. doi:10.1038/s41388-018-0541-4

Nanni, P., Landuzzi, L., Manara, M. C., Righi, A., Nicoletti, G., Cristalli, C., … Scotlandi, K. (2019). Bone sarcoma patient-derived xenografts are faithful and stable preclinical models for molecular and therapeutic investigations. Sci Rep, 9(1), 12174. doi:10.1038/s41598-019-48634-y

Nayak, R. C., Hegde, S., Althoff, M. J., Wellendorf, A. M., Mohmoud, F., Perentesis, J., … Cancelas, J. A. (2019). The signaling axis atypical protein kinase C lambda/iota-Satb2 mediates leukemic transformation of B-cell progenitors. Nat Commun, 10(1), 46. doi:10.1038/s41467-018-07846-y

Nazarian, R., Shi, H., Wang, Q., Kong, X., Koya, R. C., Lee, H., … Lo, R. S. (2010). Melanomas acquire resistance to B-RAF(V600E) inhibition by RTK or N-RAS upregulation. Nature, 468(7326), 973–977. doi:10.1038/nature09626

Nieto, M. A., Huang, R. Y., Jackson, R. A., & Thiery, J. P. (2016). Emt: 2016. Cell, 166(1), 21–45. doi:10.1016/j.cell.2016.06.028

Okuno, H., Renault Mihara, F., Ohta, S., Fukuda, K., Kurosawa, K., Akamatsu, W., … Okano, H. (2017). CHARGE syndrome modeling using patient-iPSCs reveals defective migration of neural crest cells harboring CHD7 mutations. Elife, 6. doi:10.7554/eLife.21114

Paz, H., Pathak, N., & Yang, J. (2014). Invading one step at a time: the role of invadopodia in tumor metastasis. Oncogene, 33(33), 4193–4202. doi:10.1038/onc.2013.393

Perez-Guijarro, E., Day, C. P., Merlino, G., & Zaidi, M. R. (2017). Genetically engineered mouse models of melanoma. Cancer, 123(S11), 2089–2103. doi:10.1002/cncr.30684

Priestley, P., Baber, J., Lolkema, M. P., Steeghs, N., de Bruijn, E., Shale, C., … Cuppen, E. (2019). Pan-cancer whole-genome analyses of metastatic solid tumours. Nature. doi:10.1038/s41586-019-1689-y

Rainger, J. K., Bhatia, S., Bengani, H., Gautier, P., Rainger, J., Pearson, M., … Fitzpatrick, D. R. (2014). Disruption of SATB2 or its long-range cis-regulation by SOX9 causes a syndromic form of Pierre Robin sequence. Hum Mol Genet, 23(10), 2569–2579. doi:10.1093/hmg/ddt647

Rambow, F., Rogiers, A., Marin-Bejar, O., Aibar, S., Femel, J., Dewaele, M., … Marine, J. C. (2018). Toward Minimal Residual Disease-Directed Therapy in Melanoma. Cell, 174(4), 843–855 e819. doi:10.1016/j.cell.2018.06.025

Rebecca, V. W., Wood, E., Fedorenko, I. V., Paraiso, K. H., Haarberg, H. E., Chen, Y., … Smalley, K. S. (2014). Evaluating melanoma drug response and therapeutic escape with quantitative proteomics. Mol Cell Proteomics, 13(7), 1844–1854. doi:10.1074/mcp.M113.037424

Savarese, F., Dávila, A., Nechanitzky, R., De La Rosa-Velazquez, I., Pereira, C. F., Engelke, R., … Grosschedl, R. (2009). Satb1 and Satb2 regulate embryonic stem cell differentiation and Nanog expression. Genes Dev, 23(22), 2625–2638. doi:10.1101/gad.1815709

Schulz, Y., Wehner, P., Opitz, L., Salinas-Riester, G., Bongers, E. M., van Ravenswaaij-Arts, C. M., … Pauli, S. (2014). CHD7, the gene mutated in CHARGE syndrome, regulates genes involved in neural crest cell guidance. Hum Genet, 133(8), 997–1009. doi:10.1007/s00439-014-1444-2

Shain, A. H., Yeh, I., Kovalyshyn, I., Sriharan, A., Talevich, E., Gagnon, A., … Bastian, B. C. (2015). The Genetic Evolution of Melanoma from Precursor Lesions. N Engl J Med, 373(20), 1926–1936. doi:10.1056/NEJMoa1502583

Sheehan-Rooney, K., Palinkasova, B., Eberhart, J. K., & Dixon, M. J. (2010). A cross-species analysis of Satb2 expression suggests deep conservation across vertebrate lineages. Dev Dyn, 239(12), 3481–3491. doi:10.1002/dvdy.22483

Sheehan-Rooney, K., Swartz, M. E., Lovely, C. B., Dixon, M. J., & Eberhart, J. K. (2013). Bmp and Shh signaling mediate the expression of satb2 in the pharyngeal arches. PLoS One, 8(3), e59533. doi:10.1371/journal.pone.0059533

Shinmyo, Y., & Kawasaki, H. (2017). CRISPR/Cas9-Mediated Gene Knockout in the Mouse Brain Using In Utero Electroporation. Curr Protoc Neurosci, 79, 3 32 31–33 32 11. doi:10.1002/cpns.26

Shirley, S. H., Greene, V. R., Duncan, L. M., Torres Cabala, C. A., Grimm, E. A., & Kusewitt, D. F. (2012). Slug expression during melanoma progression. Am J Pathol, 180(6), 2479–2489. doi:10.1016/j.ajpath.2012.02.014

Snyder, A., Makarov, V., Merghoub, T., Yuan, J., Zaretsky, J. M., Desrichard, A., … Chan, T. A. (2014). Genetic basis for clinical response to CTLA-4 blockade in melanoma. N Engl J Med, 371(23), 2189–2199. doi:10.1056/NEJMoa1406498

Tan, J. L., Fogley, R. D., Flynn, R. A., Ablain, J., Yang, S., Saint-Andre, V., … Zon, L. I. (2016). Stress from Nucleotide Depletion Activates the Transcriptional Regulator HEXIM1 to Suppress Melanoma. Mol Cell, 62(1), 34–46. doi:10.1016/j.molcel.2016.03.013

Tirosh, I., Izar, B., Prakadan, S. M., Wadsworth, M. H., 2nd, Treacy, D., Trombetta, J. J., … Garraway, L. A. (2016). Dissecting the multicellular ecosystem of metastatic melanoma by single-cell RNA-seq. Science, 352(6282), 189–196. doi:10.1126/science.aad0501

Van Allen, E. M., Miao, D., Schilling, B., Shukla, S. A., Blank, C., Zimmer, L., … Garraway, L. A. (2015). Genomic correlates of response to CTLA-4 blockade in metastatic melanoma. Science, 350(6257), 207–211. doi:10.1126/science.aad0095

Van Allen, E. M., Wagle, N., Sucker, A., Treacy, D. J., Johannessen, C. M., Goetz, E. M., … Dermatologic Cooperative Oncology Group of, G. (2014). The genetic landscape of clinical resistance to RAF inhibition in metastatic melanoma. Cancer Discov, 4(1), 94–109. doi:10.1158/2159-8290.CD-13-0617

van Rooijen, E., Fazio, M., & Zon, L. I. (2017). From fish bowl to bedside: The power of zebrafish to unravel melanoma pathogenesis and discover new therapeutics. Pigment Cell Melanoma Res, 30(4), 402–412. doi:10.1111/pcmr.12592

Venkatesan, A. M., Vyas, R., Gramann, A. K., Dresser, K., Gujja, S., Bhatnagar, S., … Ceol, C. J. (2018). Ligand-activated BMP signaling inhibits cell differentiation and death to promote melanoma. The Journal of clinical investigation, 128(1), 294–308. doi:10.1172/JCI92513

Verfaillie, A., Imrichova, H., Atak, Z. K., Dewaele, M., Rambow, F., Hulselmans, G., … Aerts, S. (2015). Decoding the regulatory landscape of melanoma reveals TEADS as regulators of the invasive cell state. Nat Commun, 6, 6683. doi:10.1038/ncomms7683

Wang, Y. Q., Jiang, D. M., Hu, S. S., Zhao, L., Wang, L., Yang, M. H., … Wang, S. (2019). SATB2-AS1 Suppresses Colorectal Carcinoma Aggressiveness by Inhibiting SATB2-Dependent Snail Transcription and Epithelial-Mesenchymal Transition. Cancer Res, 79(14), 3542–3556. doi:10.1158/0008-5472.CAN-18-2900

White, R. M., Sessa, A., Burke, C., Bowman, T., LeBlanc, J., Ceol, C., … Zon, L. I. (2008). Transparent adult zebrafish as a tool for in vivo transplantation analysis. Cell Stem Cell, 2(2), 183–189. doi:10.1016/j.stem.2007.11.002

Wu, F., Jordan, A., Kluz, T., Shen, S., Sun, H., Cartularo, L. A., & Costa, M. (2016). SATB2 expression increased anchorage-independent growth and cell migration in human bronchial epithelial cells. Toxicol Appl Pharmacol, 293, 30–36. doi:10.1016/j.taap.2016.01.008

Xu, H. Y., Fang, W., Huang, Z. W., Lu, J. C., Wang, Y. Q., Tang, Q. L., … Wang, J. (2017). Metformin reduces SATB2-mediated osteosarcoma stem cell-like phenotype and tumor growth via inhibition of N-cadherin/NF-kB signaling. Eur Rev Med Pharmacol Sci, 21(20), 4516–4528. Retrieved from https://www.ncbi.nlm.nih.gov/pubmed/29131265

Yen, J., White, R. M., Wedge, D. C., Van Loo, P., de Ridder, J., Capper, A., … Futreal, P. A. (2013). The genetic heterogeneity and mutational burden of engineered melanomas in zebrafish models. Genome Biol, 14(10), R113. doi:10.1186/gb-2013-14-10-r113

Yu, W., Ma, Y., Ochoa, A. C., Shankar, S., & Srivastava, R. K. (2017). Cellular transformation of human mammary epithelial cells by SATB2. Stem Cell Res, 19, 139–147. doi:10.1016/j.scr.2017.01.011

Yu, W., Ma, Y., Shankar, S., & Srivastava, R. K. (2017). SATB2/beta-catenin/TCF-LEF pathway induces cellular transformation by generating cancer stem cells in colorectal cancer. Sci Rep, 7(1), 10939. doi:10.1038/s41598-017-05458-y

Zarate, Y. A., Bosanko, K. A., Caffrey, A. R., Bernstein, J. A., Martin, D. M., Williams, M. S., … Fish, J. L. (2019). Mutation update for the SATB2 gene. Hum Mutat, 40(8), 1013–1029. doi:10.1002/humu.23771

Zarate, Y. A., & Fish, J. L. (2017). SATB2-associated syndrome: Mechanisms, phenotype, and practical recommendations. Am J Med Genet A, 173(2), 327–337. doi:10.1002/ajmg.a.38022

Zhou, L. Q., Wu, J., Wang, W. T., Yu, W., Zhao, G. N., Zhang, P., … Liu, D. P. (2012). The AT-rich DNA-binding protein SATB2 promotes expression and physical association of human (G)gamma-and (A)gamma-globin genes. J Biol Chem, 287(36), 30641–30652. doi:10.1074/jbc.M112.355271

Zhou, P., Wu, G., Zhang, P., Xu, R., Ge, J., Fu, Y., … Jiang, H. (2016). SATB2-Nanog axis links age-related intrinsic changes of mesenchymal stem cells from craniofacial bone. Aging (Albany NY), 8(9), 2006–2011. doi:10.18632/aging.101041

## Experimental Methods references

Ablain, J., Durand, E.M., Yang, S., Zhou, Y., and Zon, L.I. (2015). A CRISPR/Cas9 vector system for tissue-specific gene disruption in zebrafish. Dev Cell 32, 756–764.

Ceol, C.J., Houvras, Y., Jane-Valbuena, J., Bilodeau, S., Orlando, D.A., Battisti, V., Fritsch, L., Lin, W.M., Hollmann, T.J., Ferre, F., et al. (2011). The histone methyltransferase SETDB1 is recurrently amplified in melanoma and accelerates its onset. Nature 471, 513–517.

Dang, M., Henderson, R.E., Garraway, L.A., and Zon, L.I. (2016). Long-term drug administration in the adult zebrafish using oral gavage for cancer preclinical studies. Dis Model Mech 9, 811–820.

Gagnon, J.A., Valen, E., Thyme, S.B., Huang, P., Akhmetova, L., Pauli, A., Montague, T.G., Zimmerman, S., Richter, C., and Schier, A.F. (2014). Efficient mutagenesis by Cas9 protein-mediated oligonucleotide insertion and large-scale assessment of single-guide RNAs. PLoS One 9, e98186.

Gubelmann, C., Gattiker, A., Massouras, A., Hens, K., David, F., Decouttere, F., Rougemont, J., and Deplancke, B. (2011). GETPrime: a gene-or transcript-specific primer database for quantitative real-time PCR. Database: the journal of biological databases and curation 2011, bar040.

Heilmann, S., Ratnakumar, K., Langdon, E., Kansler, E., Kim, I., Campbell, N.R., Perry, E., McMahon, A., Kaufman, C., van Rooijen, E., et al. (2015). A quantitative system for studying metastasis using transparent zebrafish. Cancer Res.

Hiller, M., Agarwal, S., Notwell, J.H., Parikh, R., Guturu, H., Wenger, A.M., and Bejerano, G. (2013). Computational methods to detect conserved non-genic elements in phylogenetically isolated genomes: application to zebrafish. Nucleic Acids Res 41, e151.

Hu, Y., and Smyth, G.K. (2009). ELDA: extreme limiting dilution analysis for comparing depleted and enriched populations in stem cell and other assays. J Immunol Methods 347, 70–78.

Hulsen, T., de Vlieg, J., and Alkema, W. (2008). BioVenn - a web application for the comparison and visualization of biological lists using area-proportional Venn diagrams. BMC genomics 9, 488.

Jao, L.E., Wente, S.R., and Chen, W. (2013). Efficient multiplex biallelic zebrafish genome editing using a CRISPR nuclease system. Proc Natl Acad Sci U S A 110, 13904–13909.

Kim, H.J., Lee, H.J., Kim, H., Cho, S.W., and Kim, J.S. (2009). Targeted genome editing in human cells with zinc finger nucleases constructed via modular assembly. Genome Res 19, 1279–1288.

Kutner, R.H., Zhang, X.Y., and Reiser, J. (2009). Production, concentration and titration of pseudotyped HIV-1-based lentiviral vectors. Nature protocols 4, 495–505.

Langmead, B., and Salzberg, S.L. (2012). Fast gapped-read alignment with Bowtie 2. Nature methods 9, 357–359.

Lee, T.I., Johnstone, S.E., and Young, R.A. (2006). Chromatin immunoprecipitation and microarray-based analysis of protein location. Nature protocols 1, 729–748.

Loven, J., Hoke, H.A., Lin, C.Y., Lau, A., Orlando, D.A., Vakoc, C.R., Bradner, J.E., Lee, T.I., and Young, R.A. (2013). Selective inhibition of tumor oncogenes by disruption of super-enhancers. Cell 153, 320–334.

Martin, K.H., Hayes, K.E., Walk, E.L., Ammer, A.G., Markwell, S.M., and Weed, S.A. (2012). Quantitative measurement of invadopodia-mediated extracellular matrix proteolysis in single and multicellular contexts. J Vis Exp, e4119.

Meerbrey, K.L., Hu, G., Kessler, J.D., Roarty, K., Li, M.Z., Fang, J.E., Herschkowitz, J.I., Burrows, A.E., Ciccia, A., Sun, T., et al. (2011). The pINDUCER lentiviral toolkit for inducible RNA interference in vitro and in vivo. Proc Natl Acad Sci U S A 108, 3665–3670.

Subramanian, A., Tamayo, P., Mootha, V.K., Mukherjee, S., Ebert, B.L., Gillette, M.A., Paulovich, A., Pomeroy, S.L., Golub, T.R., Lander, E.S., et al. (2005). Gene set enrichment analysis: a knowledge-based approach for interpreting genome-wide expression profiles. Proceedings of the National Academy of Sciences of the United States of America 102, 15545–15550.

Whyte, W.A., Orlando, D.A., Hnisz, D., Abraham, B.J., Lin, C.Y., Kagey, M.H., Rahl, P.B., Lee, T.I., and Young, R.A. (2013). Master transcription factors and mediator establish super-enhancers at key cell identity genes. Cell 153, 307–319.

Zhang, Y., Liu, T., Meyer, C.A., Eeckhoute, J., Johnson, D.S., Bernstein, B.E., Nusbaum, C., Myers, R.M., Brown, M., Li, W., et al. (2008). Model-based analysis of ChIP-Seq (MACS). Genome biology 9, R137.

## References

Fu, J., Li, K., Zhang, W., Wan, C., Zhang, J., Jiang, P., & Liu, X. S. (2020). Large-scale public data reuse to model immunotherapy response and resistance. Genome medicine, 12(1), 21. doi:10.1186/s13073-020-0721-z

Gao, J., Aksoy, B. A., Dogrusoz, U., Dresdner, G., Gross, B., Sumer, S. O., … Schultz, N. (2013). Integrative analysis of complex cancer genomics and clinical profiles using the cBioPortal. Sci Signal, 6(269), pl1. doi:10.1126/scisignal.2004088

